# A Ca²⁺-mediated signaling pathway modulates ion homeostasis via HKT1;1 during Arabidopsis seed germination under salt stress

**DOI:** 10.64898/2026.01.25.701549

**Authors:** Yvonne Kiere, Ancy E.J. Chandran, Guy Sobol, Omer Cremer, Shaked Azoulay-Portal, Yonatan Wexler, Doron Shkolnik

## Abstract

Soil salinization severely constrains seedling establishment by disrupting cellular Na⁺/K⁺ homeostasis. Calcium (Ca²⁺) signaling contributes to the re-establishment of ion balance during early growth, yet the mechanisms linking Ca²⁺ perception to ion transport regulation remain unclear. Here, we identify a Ca²⁺-responsive regulatory module in Arabidopsis thaliana comprising CALMODULIN-BINDING TRANSCRIPTION ACTIVATOR 6 (CAMTA6), the TYPE 2C PROTEIN PHOSPHATASE PP2C49, and the HIGH-AFFINITY K⁺ TRANSPORTER HKT1;1, and define their coordinated roles during germination under salt stress. Spatial promoter analyses revealed that NaCl induces *CAMTA6* expression at cotyledon margins, while CaCl₂ stimulates *HKT1;1* transcription in the radicle, consistent with CAMTA6-mediated repression of *HKT1;1*. In *camta6* mutants, *PP2C49* expression expanded beyond its normal radicle-restricted domain, indicating CAMTA6-dependent spatial control. Promoter activation assays *in planta* demonstrated CAMTA6-dependent transactivation of the *HKT1;1* and *PP2C49* promoters. Treatment with the PP2C inhibitor sanguinarine enhanced germination under salinity in the wild type, but not in *hkt1* nor in the salt-tolerant *camta6* and *pp2c49* mutants. Sanguinarine restricted *CAMTA6* promoter activity to cotyledon margins, suppressed *PP2C49* expression, and enhanced *HKT1;1* accumulation in the radicle, collectively supporting improved Na⁺/K⁺ balance. Transcriptome profiling further revealed additional Ca²⁺-responsive PP2C genes under CAMTA6-dependent regulation. Together, these findings establish a Ca²⁺-regulated transcriptional network coordinating ion homeostasis during germination and suggest strategies to support seedling performance in saline environments.

**Significance statement:** Salinity impairs germination largely by disrupting Na⁺/K⁺ homeostasis, yet the signaling pathways that protect seedlings at this stage remain poorly defined. We identify a calcium-responsive regulatory mechanism that spatially coordinates transcriptional control of genes involved in ion transport during early development, providing a mechanistic basis for improving seedling establishment in saline soils.

## INTRODUCTION

Salt stress is a major abiotic factor limiting plant growth and productivity, particularly during seed germination and early seedling establishment (Munns & Tester, 2008). Increasing salinization of agricultural soils worldwide (Wang et al., 2003; Hassani et al., 2021) highlights the need for a deeper understanding of the mechanisms governing plant responses to salinity, particularly during the early stages of development. During seed germination, salt stress causes osmotic stress, disrupting water uptake and seed imbibition; and oxidative stress, leading to the accumulation of reactive oxygen species that damage cellular structures. In addition, high salinity results in ion imbalances, primarily through excessive accumulation of sodium (Na⁺), which impairs metabolic processes that are necessary for germination and seedling growth (Zhu, 2002; Munns and Tester, 2008).

In contrast to Na^+^, potassium (K⁺) is an essential macronutrient that plays a vital role in various physiological processes during seed germination, including enzyme activation, osmoregulation, and maintenance of cellular turgor. Under salt stress, elevated Na⁺ levels can disrupt K⁺ homeostasis, leading to reduced K⁺ uptake and accumulation in germinating seedlings. This imbalance adversely affects germination rates and seedling vigor (Wang et al., 2013; Wang and Wu, 2013; Sun et al., 2015).

Plants employ various regulatory mechanisms to maintain ion homeostasis under saline conditions. One major strategy is to reduce harmful Na⁺ accumulation in shoot tissues by retrieving Na⁺ from the root xylem stream, a process mediated by Na⁺/K⁺ transporters (HKTs) (Uozumi et al., 2000; Mäser et al., 2002; Assaha et al., 2017). This shoot-protective activity from salinity has been demonstrated in the model plant *Arabidopsis thaliana*, as well as in important crop plants, including wheat (*Triticum aestivum*) and rice (*Oryza sativa*) (Hauser and Horie, 2010; Byrt et al., 2014). In Arabidopsis, HKT1;1 is the only HKT family member, playing a pivotal role in controlling whole-plant Na^+^/K^+^ homeostasis and ensuring proper seedling (Mäser et al., 2002; Berthomieu et al., 2003; Horie et al., 2006; Davenport et al., 2007). In germinating Arabidopsis seedlings, HKT1;1 is negatively regulated at the transcriptional level by calcium (Ca²⁺) via CALMODULIN-BINDING TRANSCRIPTION ACTIVATOR 6 (CAMTA6) (Shkolnik et al., 2019; Chandran et al., 2024), and post-translationally by the type 2C protein phosphatase PP2C49 (Chu et al., 2021). In the *camta6* mutant background, expression of *HKT1;1* is confined to the radicle tissues—an expression pattern associated with improved germination rates under high Na⁺ concentrations. This mutant’s enhanced salt tolerance is likely achieved by maintaining low Na⁺/K⁺ ratios in the germinating seedling (Shkolnik et al., 2019; Chandran et al., 2024). In this study, we further elucidate the regulatory interplay between Ca²⁺ signaling, CAMTA6, PP2C49, and HKT1;1 in controlling *Arabidopsis* seed germination under salt-stress conditions.

A key component of the plant’s adaptive response involves calcium-mediated signaling pathways that decode salt-induced cytosolic Ca²⁺ transients and modulate gene expression and hormone sensitivity (Dodd et al., 2010). Calcium sensors such as calcineurin B-like proteins (CBLs), CBL-interacting protein kinases, calmodulin-like proteins, and calcium-dependent protein kinases integrate salt-stress signals with abscisic acid (ABA) and gibberellin pathways to inhibit germination(Pandey et al., 2004; Yip Delormel and Boudsocq, 2019). This tightly regulated signaling network ensures that germination proceeds only under favorable conditions.

CAMTAs are key regulators of plant responses to environmental stress, integrating calcium signaling with gene expression. In Arabidopsis, the CAMTA family consists of six members (CAMTA1–CAMTA6), which participate in diverse physiological processes, including adaptation to abiotic and biotic stresses (Finkler et al., 2007a; Galon et al., 2010; Rahman et al., 2016). CAMTAs are known to mediate plant responses to cold, drought, and salinity by modulating stress-responsive gene expression, particularly through interactions with calcium/calmodulin signaling cascades (Yang et al., 2012; Kim et al., 2017). Among them, CAMTA6 has emerged as a crucial player in salt-stress adaptation, particularly during seed germination, where it functions as a negative regulator of salt tolerance (Shkolnik et al., 2019). Mutants lacking functional CAMTA6 exhibit enhanced germination rates under high-salt conditions, suggesting that the absence of CAMTA6 enhances seed germination by improving Na^+^/K^+^ homeostasis (Chandran et al., 2024). Understanding CAMTA6-mediated transcriptional networks can offer valuable insights into the regulation of germination under challenging environmental conditions.

Protein phosphatase 2C (PP2C) proteins play crucial roles in abiotic stress signaling, including responses to salt stress during seed germination (Schweighofer et al., 2004; Park et al., 2009; Chandran et al., 2024). In the Arabidopsis genome, more than 80 *PP2C* genes have been identified. Based on sequence similarity and domain organization, these *PP2C*s are classified into 10 distinct groups, designated A through J (Schweighofer et al., 2004). Group A PP2Cs, including ABI1, ABI2, HAB1, and HAB2, function as negative regulators of ABA signaling, a key pathway involved in stress responses (Fujii et al., 2009). These phosphatases dephosphorylate and inactivate SNF1-related protein kinase 2 enzymes, thereby suppressing ABA-mediated stress adaptation. During seed germination under salt stress, ABA levels increase to germination-arresting concentrations, and PP2Cs play a crucial role in modulating the balance between germination and dormancy by controlling ABA sensitivity (Park et al., 2009; Antoni et al., 2012). PP2C49 is a group G PP2C that has been implicated in the dephosphorylation of the sodium transporter HKT1;1, thereby regulating ion fluxes and contributing to the maintenance of cellular ion homeostasis under salt stress (Shkolnik et al., 2019; Chu et al., 2021; Chandran et al., 2024). Understanding the role of PP2Cs in salt-stress responses—whether through Ca²⁺-dependent or independent pathways—may provide valuable insights into plant-resilience mechanisms and identify potential targets for enhancing salt tolerance.

Given the growing challenges posed by global climate change and the progressive salinization of agricultural soils, understanding the molecular, cellular, and physiological mechanisms that regulate plant responses to salinity during germination is crucial. Such insights are essential for guiding breeding strategies and biotechnological interventions aimed at improving salt tolerance in crop species.

## RESULTS

### Calcium-mediated modulation of CAMTA6, HKT1;1, and PP2C49 expression during salt stress in germinating Arabidopsis

In germinating Arabidopsis seedlings, CAMTA6 is predominantly expressed in the cotyledons, whereas HKT1;1 is expressed in both the cotyledons and the radicle (Shkolnik et al., 2019; Chandran et al., 2024). We previously reported that CaCl₂ treatment induces radicle-enriched HKT1;1 expression, correlating with enhanced salt tolerance during germination (Chandran et al., 2024). To further explore how calcium and salt stress affect the spatial expression of these genes, we examined *CAMTA6* and *HKT1;1* promoter activity in transgenic lines expressing GUS under the control of each promoter (*CAMTA6pro:GUS* and *HKT1;1pro:GUS*) (Shkolnik et al., 2019). Under all tested conditions, *HKT1;1* promoter activity was absent from the cotyledon margins. In contrast, *CAMTA6* expression became focused or markedly enhanced in these margins following CaCl₂ or NaCl treatment (Figure. 1a). Notably, under combined NaCl and CaCl₂ treatment, *CAMTA6* expression was strictly confined to the cotyledon margins, whereas *HKT1;1* expression was mainly restricted to the radicle (Figure 1a), indicating a spatially segregated and stress-dependent transcriptional response.

**Figure 1.**
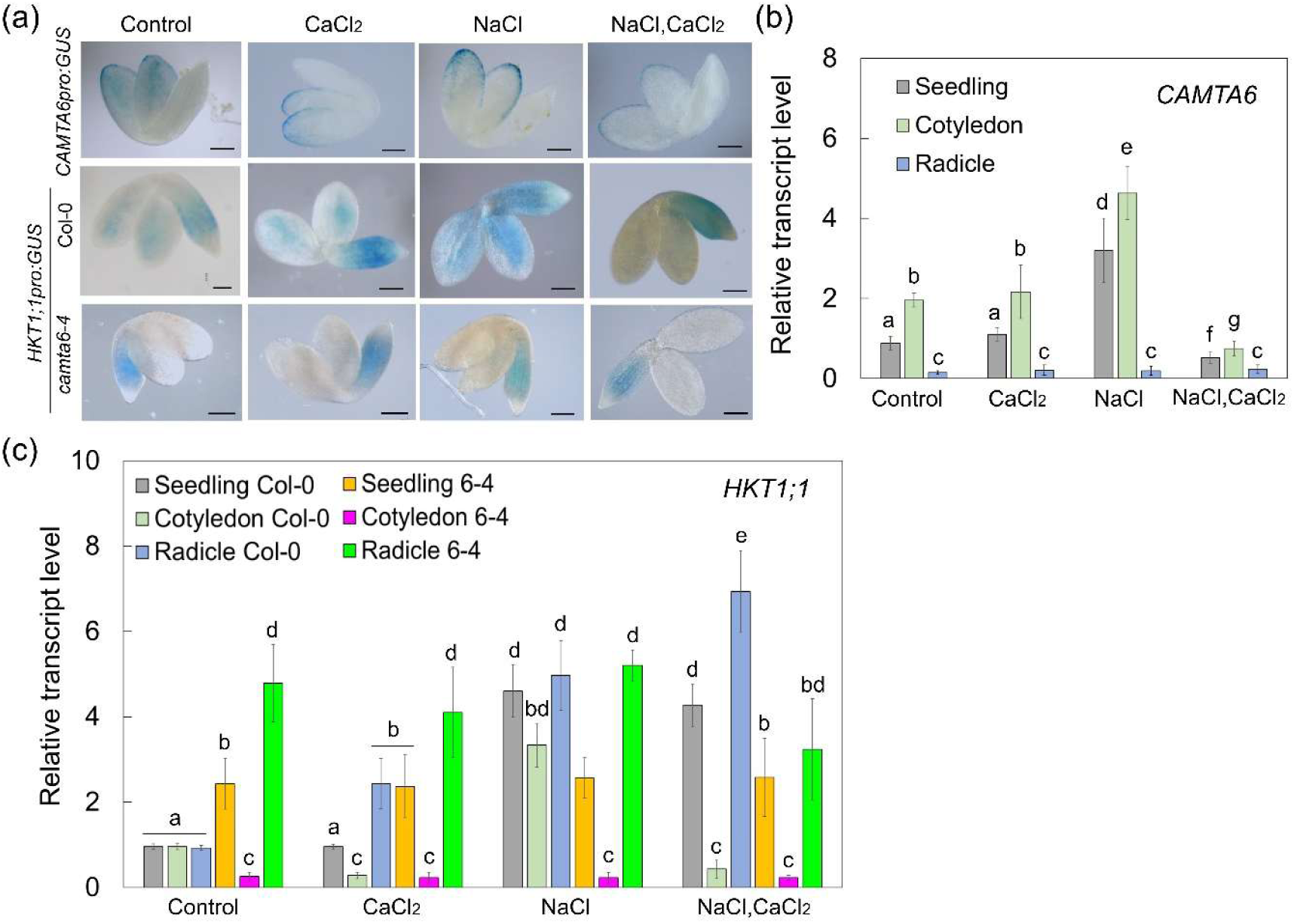
NaCl and CaCl₂ modulate CAMTA6 and HKT1;1 expression in wild-type and *camta6-4* Arabidopsis seedlings. (a) GUS staining of seedlings harboring *CAMTA6pro:GUS* or *HKT1;1pro:GUS* and treated with single or combined chemicals, as indicated (NaCl 150 mM; CaCl₂ 10 mM) for 16 h before ejection, staining, and imaging (bars = 0.5 mm). (b, c) RT-qPCR quantification of relative transcript levels of CAMTA6 (b) and HKT1;1 (c) in pre-germinated seedlings. Data are presented as means ± SD from three independent biological experiments (∼30 seeds each). Different lowercase letters indicate significantly different values by Tukey’s HSD post hoc test (P < 0.01).

Similarly, in the *camta6-4* mutant background, *HKT1;1* expression was confined to the radicle (Figure 1a), indicating that both the mutation and the combined CaCl₂–NaCl treatment elicit comparable spatial regulation of this gene. To validate the spatial expression patterns of *CAMTA6* and *HKT1;1*, we performed reverse transcription quantitative PCR (RT-qPCR) on RNA extracted from whole seedlings, cotyledons, and radicles (see Materials and Methods). Consistent with the promoter activity data, *CAMTA6* transcripts were predominantly detected in cotyledons, with substantially lower expression in radicles under all treatments (Figure 1B). Upon NaCl exposure, *CAMTA6* expression in the cotyledons increased substantially (4.63 ± 0.06, relative to whole seedlings set to 1.0), whereas expression in the radicle remained low (0.2 ± 0.1) (Figure 1b). Under the combined CaCl₂–NaCl treatment, cotyledon *CAMTA6* transcript levels (1.52 ± 0.18) were reduced relative to CaCl₂ alone (2.16 ± 0.15) and untreated controls (1.96 ± 0.18) (Figure 1b). Under the same combined treatment, *HKT1;1* transcript was strongly enriched in the radicle (6.93 ± 0.96) and low in the cotyledons (0.43 ± 0.2) (Figure 1c). In the *camta6-4* mutant, *HKT1;1* expression remained confined to the radicle under all tested conditions, consistent with the promoter activity results (Figure 1, a and c). Together, these findings suggest that Ca²⁺ promotes spatially opposite expression gradients for *CAMTA6* and *HKT1;1*, a pattern that becomes more pronounced under salt stress.

*HKT1;1* is regulated transcriptionally by CAMTA6 (Shkolnik et al., 2019) and post-translationally inactivated by PP2C49 through direct dephosphorylation (Chu et al., 2021). We recently showed that *PP2C49* is expressed in the radicle and that CaCl₂ treatment suppresses its NaCl-induced expression (Chandran et al., 2024) (Figure 2). To investigate whether CAMTA6 also regulates *PP2C49* at the transcriptional level, we analyzed *PP2C49* promoter activity in *camta6-4* seedlings expressing the *PP2C49pro:GUS* reporter (Chandran et al., 2024). Remarkably, in the *camta6-4* background, *PP2C49* promoter activity was strongly upregulated in all seedling tissues under all tested conditions (Figure 2a). This widespread expression pattern was corroborated by RT-qPCR analysis (Figure 2b). These findings suggest that CAMTA6 plays a central role in repressing *PP2C49* transcription during germination, thereby potentially contributing to the spatial control of HKT1;1 activity under salt stress.

**Figure 2.**
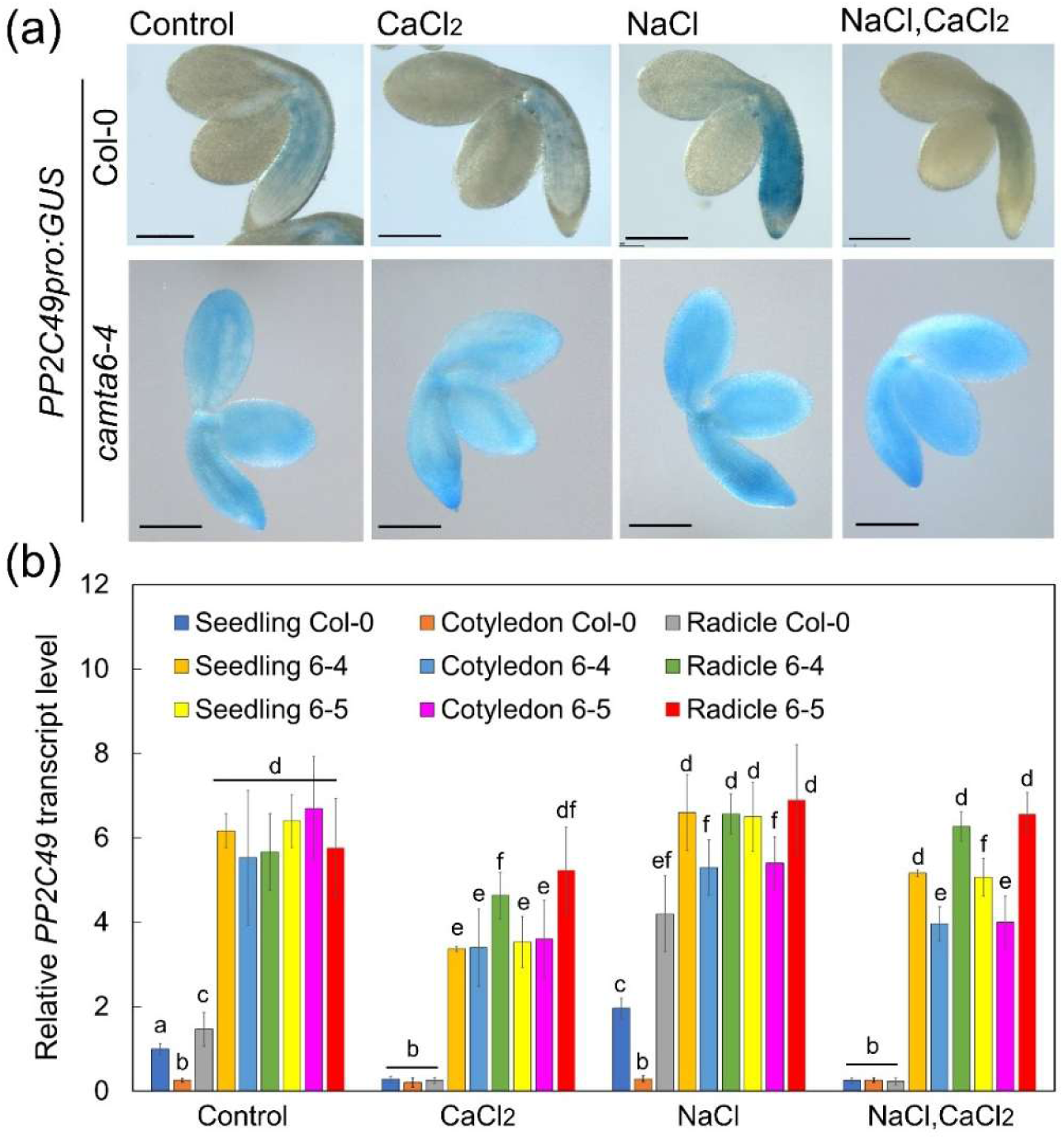
NaCl and CaCl₂ regulate *PP2C49* expression in wild-type and *camta6-4* mutant Arabidopsis seedlings. (a) GUS staining of ejected seedlings harboring *PP2C49pro:GUS* and treated with single or combined chemicals as indicated (NaCl 150 mM; CaCl₂ 10 mM) for 16 h before ejection, staining, and imaging (bars = 0.5 mm). (b) RT-qPCR quantification of relative *PP2C49* transcript levels in pre-germinated seedlings. Data are presented as means ± SD from three independent biological experiments (∼30 seeds each). Different lowercase letters indicate significantly different values by Tukey’s HSD post hoc test (*P* < 0.01).

At later developmental stages (4- to 11-day-old seedlings), *camta6* mutants exhibit hypersensitivity to salt stress (Shkolnik et al., 2019). To further explore the roles of CAMTA6, PP2C49, and HKT1;1 in the Ca²⁺-mediated response to NaCl, we examined their expression patterns in 5-day-old whole seedlings and their primary root tips following treatment with CaCl₂, NaCl, or both. All three genes were expressed in the root vasculature, with enhanced promoter activity observed at the hypocotyl–root junction (Figure S1). *CAMTA6* expression was induced by each treatment—CaCl₂, NaCl, and their combination, whereas *HKT1;1* expression was only induced by the combined treatment, and this induction was absent in the *camta6-4* background (Figure S1). *PP2C49* expression was induced by NaCl, but this induction was suppressed by CaCl₂, which counteracted the NaCl effect. In the *camta6-4* mutant, *PP2C49* promoter activity was highly responsive to NaCl, and the antagonistic effect of CaCl₂ was retained (Figure S1). Notably, *PP2C49* was also expressed in the root tip columella cells, where its expression was induced by CaCl₂, NaCl, and their combination, as previously observed during germination (Figure 2a; Figure S1b). These findings suggest that Ca²⁺-dependent regulation of *HKT1;1* and *PP2C49* persists beyond germination, but displays distinct spatial and conditional dynamics at later developmental stages, as further discussed below.

CAMTA transcription factors localize to the nucleus and bind specific *cis*-elements in target gene promoters, including the ABRE/CAMTA-binding motif (CACGTG[C/T/G]) and its coupling element ABRE-CE ([C/A]ACGCG[T/C/G]) (Kaplan et al., 2006; Whalley and Knight, 2013). In *HKT1;1*, mutation of an ABRE element (ACGTGT) located 75 bp upstream of the translation start codon abolishes its promoter activity (Shkolnik et al., 2019). An additional ABRE motif was identified at position –283 bp (Figure S2). However, direct binding of CAMTA6 to the *HKT1;1* promoter has not yet been demonstrated.

We therefore screened the PP2C49 promoter for similar cis-elements and identified two canonical ABRE/CAMTA-binding motifs (CACGTGTC) at –388 and –287 bp, as well as a full ABRE-CE (CACGCGGC) at –203 bp relative to the translation start codon (Figure S2). To assess direct binding, we performed a promoter-activation assay by transiently co-expressing the GUS reporter gene driven by the HKT1;1 or PP2C49 promoter, along with the CAMTA6 DNA-binding domain (CAMTA6_CG-1_) (da Costa Silva, 1994) in *Nicotiana benthamiana* leaves (Figure S3). Constitutive expression of CAMTA6_CG-1_ led to enhanced GUS activity driven by both promoters (Figure S3a), supporting the notion that CAMTA6 can transcriptionally activate these genes. CAMTA6_CG-1_ expression was verified by RT-qPCR (Figure S3b). As a control, we tested the promoter of PP2CG1 (AT2G33700), an ABA-regulated phosphatase that enhances salt tolerance in young Arabidopsis seedlings (Liu et al., 2012), and observed no activation by CAMTA6_CG-1_ (Figure S3a,e). Although the PP2CG1 promoter contains two ABRE/CAMTA motifs at –889 and –1089 bp, their distal location may render them ineffective for CAMTA6-mediated regulation (Figure S2). Importantly, CAMTA6_CG-1_ contains only the DNA-binding and recognition domain of CAMTA6, but lacks additional regions required for transcriptional repression. Therefore, this assay selectively reports promoter activation resulting from direct CAMTA6 binding to *cis*-elements, rather than the full regulatory behavior of the native CAMTA6 protein, and only increased GUS activity was expected and observed.

### Sanguinarine treatment enhances seed germination under salt-stress conditions

PP2C49 acts as a key mediator of the Ca²⁺/CAMTA6-signaling pathway that modulates HKT1;1 activity, which in turn enhances K⁺ retention in germinating seedlings under salt stress (Shkolnik et al., 2019; Chu et al., 2021; Chandran et al., 2024). The plant-derived alkaloid sanguinarine is a selective, cell-active inhibitor of mitogen-activated protein kinase phosphatase-1 (Vogt *et al*., 2005) and has also been reported to specifically and potently inhibit PP2Cs in human cells (Aburai *et al*., 2010). Based on this, we examined the effect of sanguinarine on seed germination under salt stress in Arabidopsis wild type and mutant lines (*camta6*, *pp2c49*, *hkt1*). Remarkably, supplementation with 1 μM sanguinarine significantly improved germination rates of wild type seeds in the presence of 150 and 200 mM NaCl, from 76 ± 5.36% and 3 ± 0.23% to 92 ± 4.21% and 37 ± 3.88%, respectively (Figure 3).

**Figure 3.**
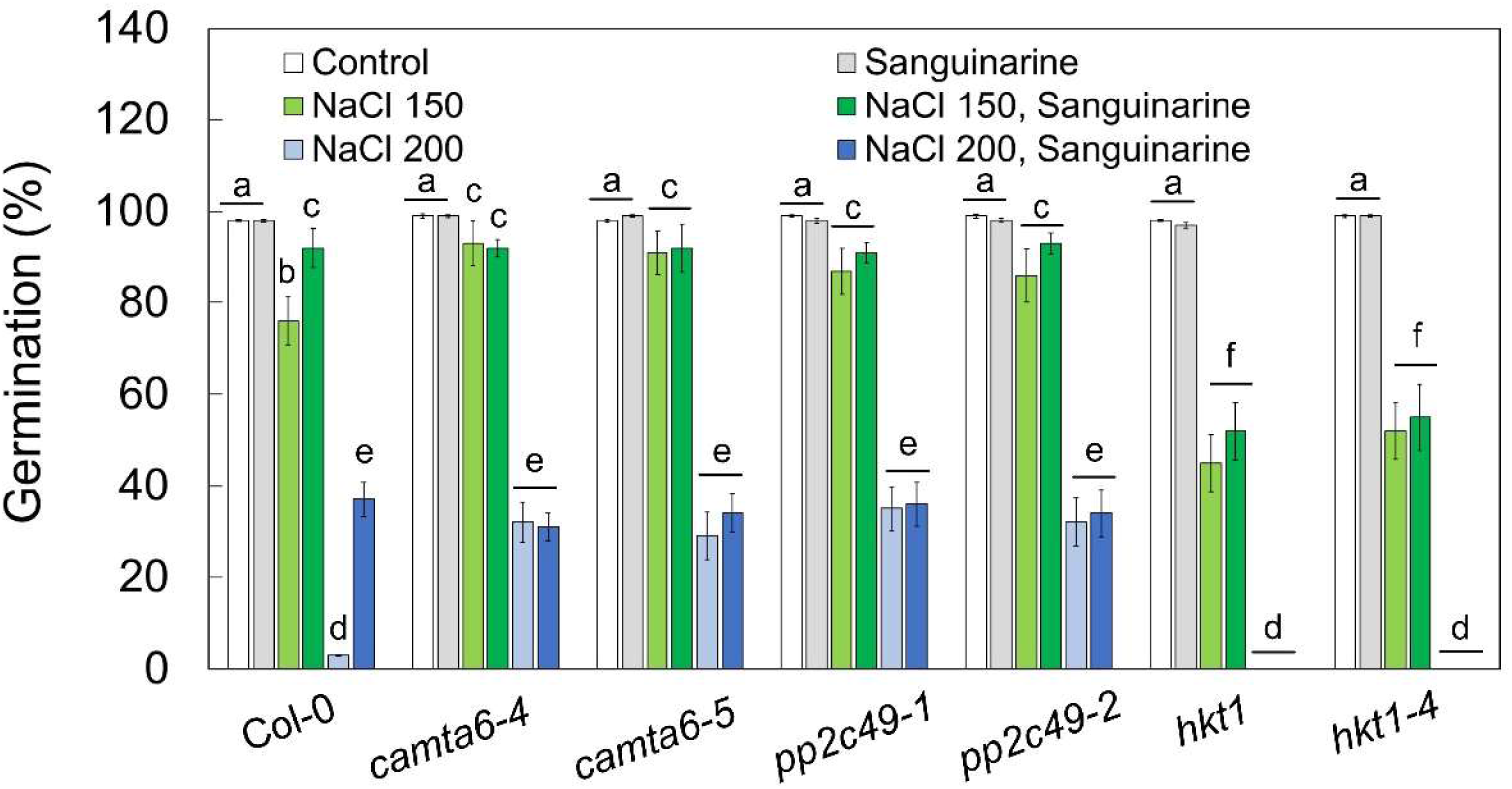
Sanguinarine enhances Arabidopsis seed germination under salt stress. Germination assay. Seeds of the indicated genotypes were sown on agar-solidified 0.25X MS medium (control) supplemented with the indicated single or combined chemicals (NaCl, 150 mM or 200 mM; sanguinarine, 1 µM). Germination rates were scored 5 days after plating. Data are presented as means ± SD from three independent biological experiments (∼100 seeds each). Different lowercase letters indicate significantly different values by Tukey’s HSD post hoc test (*P* < 0.001).

In contrast to its stimulatory effect on wild type seeds under salt stress, sanguinarine had no effect on germination rates of the salt-tolerant *camta6* and *pp2c49* mutants, or the salt-sensitive *hkt1* mutants (Shkolnik et al., 2019; Chandran et al., 2024), under identical conditions. Under control conditions, or in the presence of sanguinarine alone, no significant phenotypic differences were observed across genotypes (Figure 3). These findings support the notion that sanguinarine promotes germination under salt stress by inhibiting PP2C49, thereby preserving HKT1;1 activity, and reinforce the involvement of CAMTA6 in this Ca²⁺-dependent regulatory module.

To examine whether sanguinarine affects group A PP2Cs, which are known to mediate ABA signaling (Umezawa et al., 2009; Antoni et al., 2012), we assessed its impact on seed germination in the presence of ABA. Wild type seeds germinated at 87.6 ± 2.8%, 0%, and 0% on medium containing 100, 500, or 1000 nM ABA, respectively. Co-application of 1 μM sanguinarine—previously shown to alleviate NaCl-induced germination arrest—had no significant effect, with germination rates of 88.2 ± 5.3%, 0%, and 0% under the same respective ABA concentrations (Figure S4). These findings indicate that sanguinarine does not relieve ABA-induced inhibition of germination, and thus likely does not target the canonical ABA-associated PP2Cs during this developmental window.

Although PP2C49 is not considered part of the canonical ABA-signaling pathway, we determined whether PP2C49 and HKT1;1 are involved in ABA-dependent salt-stress responses during germination. Motivated by the enhanced germination of *camta6* mutants on ABA-containing media, we examined the germination of two allelic *pp2c49* mutants and two *hkt1* mutants under increasing concentrations of ABA. At 100 nM ABA, all genotypes germinated at ∼95%, whereas at 500 and 1000 nM ABA, germination was completely inhibited in both mutants, similar to Col-0 (Figure S5). These results indicate that *hkt1* and *pp2c49* do not exhibit altered ABA sensitivity during germination, suggesting that their salt-related phenotypes may be ABA-independent.

To further explore the role of group G PP2Cs in seed germination under salt stress, we also examined PP2CG1, a phosphatase phylogenetically related to PP2C49. Seeds of *pp2cg1* mutants did not exhibit salt tolerance comparable to that of *pp2c49* mutants. Furthermore, sanguinarine treatment enhanced the germination of *pp2cg1* mutants under salt stress, similar to its effect on wild type seeds (Figure 3 and Figure S6). These findings tentatively suggest some specificity of sanguinarine toward PP2C49, although effects on other phosphatases cannot be excluded.

### Modulation of CAMTA6, PP2C49, and HKT1;1 expression by sanguinarine during salt-stressed germination

Sanguinarine has been reported to inhibit specific PP2C-type phosphatases in mammalian systems (Aburai et al., 2010), suggesting that PP2Cs may represent a potential molecular target. Here, we will directly test whether this inhibitory relationship is conserved in plants.

Because PP2C49 is involved in regulating ion homeostasis during germination (Chandran et al., 2024), we considered whether chemical inhibition of PP2C activity might influence the seedling response to salt. Sanguinarine, a benzophenanthridine alkaloid, has been characterized in mammalian systems as a PP2C inhibitor (Aburai et al., 2010). However, its molecular function in plant cells remains largely uncharacterized. To investigate the mechanistic basis of sanguinarine’s alleviation of salt-induced germination arrest within the CAMTA6–PP2C49–HKT1;1 regulatory module, we examined the expression patterns of these genes in germinating seedlings treated with NaCl and/or sanguinarine. Notably, sanguinarine markedly reduced CAMTA6 promoter activity which, under control conditions, is broadly expressed across the cotyledon, but was restricted to the margins following treatment (Figure 1a and Figure 4a). Moreover, the NaCl-induced upregulation of CAMTA6 expression was abolished by sanguinarine treatment (Figure 1a and Figure 4a).

**Figure 4.**
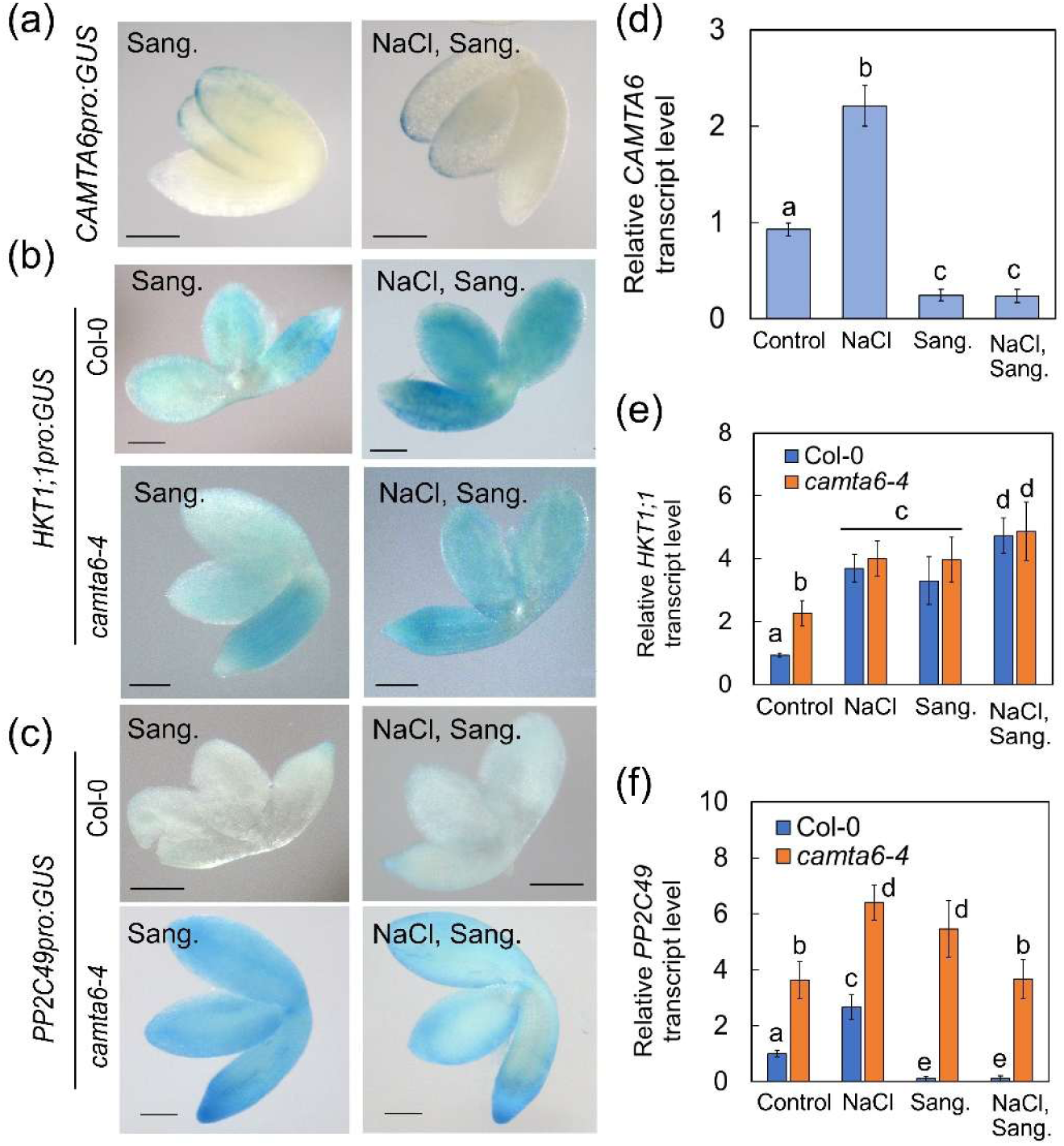
Sanguinarine and NaCl influence *CAMTA6*, *PP2C49*, and *HKT1;1* expression in germinating Arabidopsis seedlings, as revealed by GUS staining and RT-qPCR. Sanguinarine (Sang.) modulates the expression of *CAMTA6*, *PP2C49*, and *HKT1;1* in germinating *Arabidopsis*. (a–c) GUS staining of ejected wild-type (Col-0) or *camta6-4* seedlings harboring the indicated constructs and treated with 150 mM NaCl and/or 1 µM sanguinarine, as indicated (bars = 0.5 mm). (d–f) RT-qPCR quantification of relative transcript levels of *CAMTA6* (d), *HKT1;1* (E), and *PP2C49* (f) in germinating seedlings. Data are means ± SD (three biological experiments, ∼30 seeds each) and different lowercase letters indicate significantly different values by Tukey’s HSD post hoc test (*P* < 0.01).

These results were corroborated by RT-qPCR analysis of whole germinating seedlings, revealing mean ± SD relative *CAMTA6* transcript levels of 0.92 ± 0.07 (control), 2.21 ± 0.21 (NaCl), 0.24 ± 0.06 (sanguinarine), and 0.23 ± 0.07 (NaCl–sanguinarine) (Figure 4d). These data suggest that sanguinarine strongly represses CAMTA6 expression during germination, even under salt stress. Interestingly, although *camta6* mutants germinate more efficiently under salt stress (Shkolnik et al., 2019), they are hypersensitive at later seedling stages, consistent with a developmental stage-dependent function of CAMTA6 (Shkolnik et al., 2019).

Unlike seed germination, sanguinarine treatment enhanced *CAMTA6* promoter activity in the root vasculature of young seedlings, resembling the response to CaCl₂, NaCl, and their combined application (Figure S7). In wild type seedlings, NaCl treatment induced the expression of both *HKT1;1* and *PP2C49*, whereas sanguinarine suppressed both transcripts. Notably, the NaCl-induced upregulation of *PP2C49*, but not *HKT1;1*, was attenuated by sanguinarine. In the *camta6-4* background, the responsiveness of the *HKT1;1* and *PP2C49* promoters to these treatments was largely similar to that observed in wild type seedlings. However, the strong NaCl-induced activation of *PP2C49* promoter in young root zones, and its repression by sanguinarine, persisted in the *camta6-4* mutant, suggesting the involvement of a CAMTA6-independent regulatory component (Figure S7). Consistently, sanguinarine treatment did not differentially affect root development in *camta6-4*, *pp2c49-1*, or *hkt1-4* mutants compared to the wild type (Figure S8).

To examine whether sanguinarine inhibits PP2C enzymatic activity in plants, we measured phosphatase activity in crude root protein extracts of 4-day-old Arabidopsis seedlings using pNPP as substrate. Remarkably, sanguinarine significantly reduced phosphatase activity in Col-0 extracts, from 1.00 ± 0.10 to 0.16 ± 0.04 (arbitrary fluorescence units, a.u.). In contrast, extracts from *pp2c49-1* roots displayed substantially lower basal phosphatase activity compared with wild type (0.33 ± 0.09 a.u.), and this residual activity was not further reduced by sanguinarine treatment (0.09 ± 0.03 a.u.) (Figure S9). Together, these data indicate that sanguinarine inhibits phosphatase activity in Arabidopsis roots, with reduced sensitivity observed in the *pp2c49-1* mutant.

### Sanguinarine enhances K^+^ accumulation in germinating seedlings under salt-stress conditions

Exposure to excess Na^+^ levels impairs K^+^ retention and is associated with arrest of the seed-germination process (Shkolnik et al., 2019; Chandran et al., 2024). Supplementing the growth medium with CaCl₂ has been shown to enhance K^+^ accumulation and improve germination rates under salt stress (Chandran et al., 2024). Compared to wild type seedlings, germinating *hkt1* mutants accumulate higher Na^+^ and lower K^+^ levels under salt stress, whereas *pp2c49* and *camta6* mutants exhibit the opposite ion profile under the same conditions (Shkolnik et al., 2019; Chandran et al., 2024) (Figure 5). Based on these findings, we investigated whether sanguinarine influences Na^+^ and K^+^ contents in germinating wild type, *camta6*, *hkt1*, and *pp2c49* mutants in the presence of NaCl.

**Figure 5.**
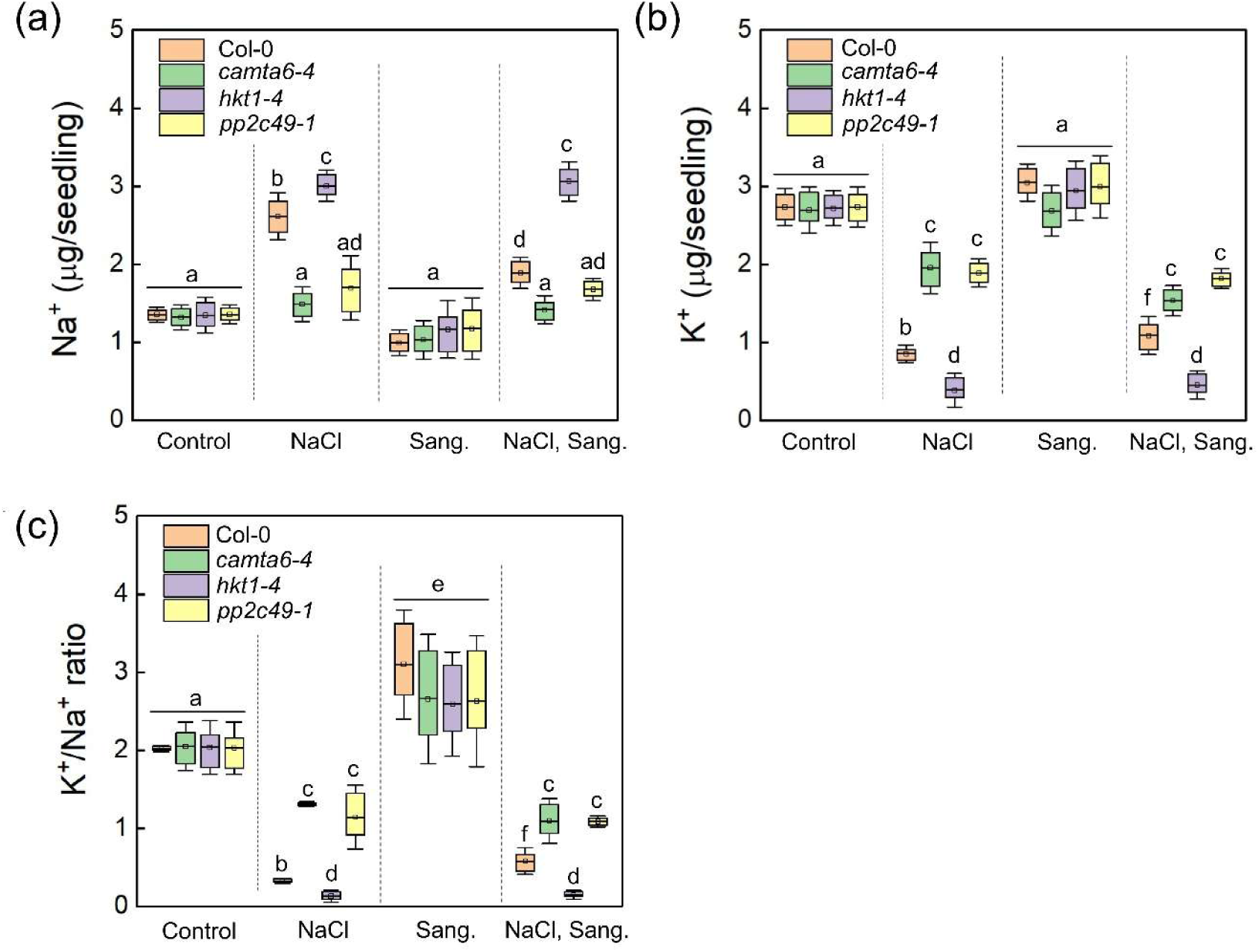
Accumulation of Na⁺ and K⁺ in germinating seedlings under NaCl and sanguinarine treatments. Accumulation of Na⁺ (a) and K⁺ (c), and K⁺/Na⁺ ratios (c) in wild type (Col-0), *camta6-4*, *pp2c49-1*, and *hkt1-4* germinating seedlings in the presence of NaCl and/or sanguinarine (Sang.). Ion content was determined by ICP-OES. Seedlings were treated with 0 (control), 1 µM Sang., 150 mM NaCl, or both, as indicated, for 16 h before seedling ejection. Data are presented as means ± SD from three independent biological experiments (∼40 seeds each). Different lowercase letters indicate significantly different values by Tukey’s HSD post hoc test (P < 0.01).

**Figure 6.**
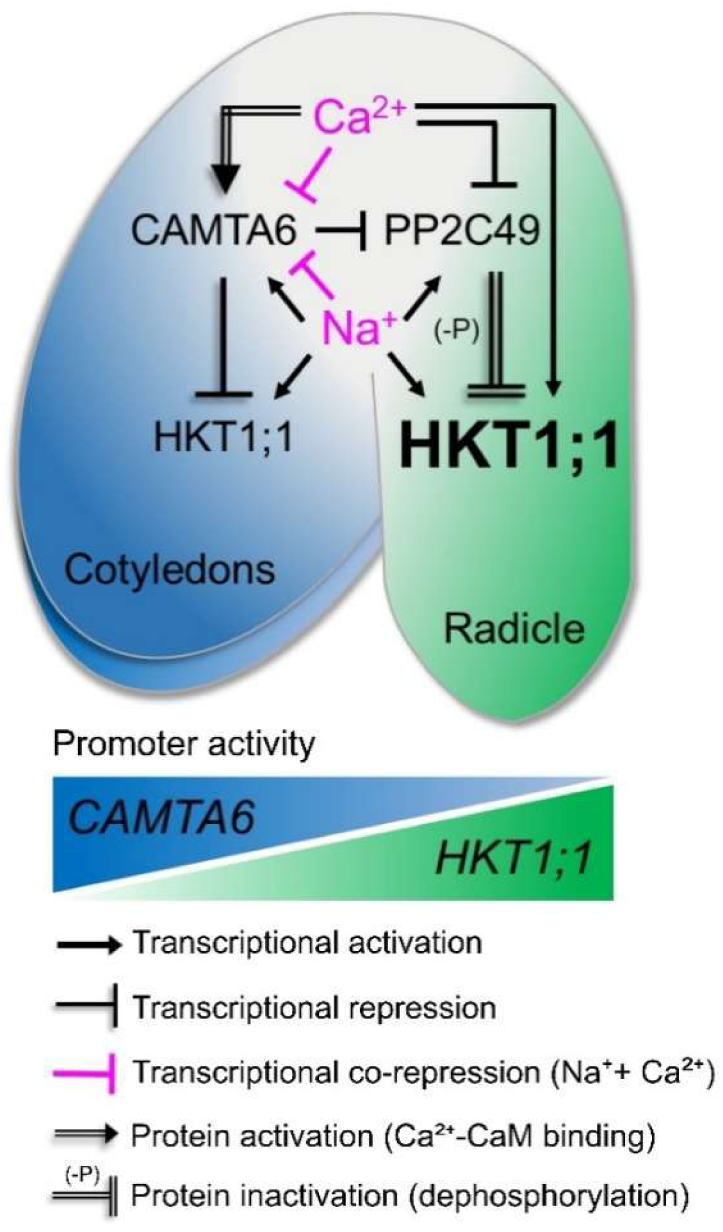
Ca²⁺-mediated signaling coordinates the expression and activity of CAMTA6, PP2C49, and HKT1;1 to integrate salt-stress responses during seed germination in this model. In this model, Ca²⁺ signaling promotes expression of the transcription factor CAMTA6 at the cotyledon margins and activates it via Ca²⁺–CaM binding, induces radicle-confined expression of the K⁺/Na⁺ symporter HKT1;1, and simultaneously represses its direct negative regulator, the phosphatase PP2C49. PP2C49 appears to inactivate HKT1;1 through direct dephosphorylation (Chu et al., 2021). In contrast, Na⁺ alone enhances CAMTA6 expression at the cotyledon margins, expands HKT1;1 expression throughout the seedling (excluding the margins), and activates PP2C49 specifically in the radicle (Fig. 1). Under combined CaCl₂ and NaCl treatment, CAMTA6 expression is attenuated (Fig. 1).

To quantify Na⁺ and K⁺ accumulation in germinating wild type, *camta6-4*, *hkt1-4*, and *pp2c49-1* seedlings, we analyzed seedling extracts using inductively coupled plasma optical emission spectrometry (ICP-OES) as previously described (Shkolnik et al., 2019). Seedlings were treated for 16 h with 1 µM sanguinarine, 150 mM NaCl, both, or no treatment (control). In response to NaCl, wild type, *camta6-4*, *hkt1-4*, and *pp2c49-1* seedlings accumulated 2.61 ± 0.2 µg, 1.49 ± 0.14 µg, 3.10 ± 0.13 µg, and 1.7 ± 0.26 µg Na⁺ per seedling, respectively (Figure 5), confirming the previously reported reduced Na⁺ accumulation in *camta6* and *pp2c49* seedlings under salt stress (Shkolnik et al., 2019; Chandran et al., 2024).

Notably, in response to the combined NaCl–sanguinarine treatment, wild type seedlings accumulated 1.89 ± 0.13 µg Na⁺ per seedling—a substantial reduction compared to NaCl treatment alone. In contrast, *hkt1-4* seedlings accumulated 3.06 ± 0.17 µg Na⁺ per seedling, similar to their response to NaCl alone (Figure 5). Similarly, *camta6-4* and *pp2c49-1* seedlings showed no significant difference in Na⁺ accumulation between NaCl and the combined treatment, with 1.41 ± 0.12 µg and 1.67 ± 0.10 µg Na⁺ per seedling, respectively (Figure 5). Under control and sanguinarine-only conditions, all genotypes accumulated comparable Na⁺ levels, approximately 1.32–1.35 µg and 0.99–1.17 µg per seedling, respectively (Figure 5).

Analysis of K⁺ concentrations in response to NaCl and NaCl–sanguinarine treatments revealed accumulation levels of 0.85 ± 0.07 µg and 1.10 ± 0.16 µg per seedling in the wild type, 1.95 ± 0.21 µg and 1.53 ± 0.13 µg in *camta6-4*, 0.38 ± 0.14 µg and 0.45 ± 0.12 µg in *hkt1-4*, and 1.90 ± 0.12 µg and 1.81 ± 0.08 µg in *pp2c49-1*, respectively (Figure 5). Under control and sanguinarine-only treatments, all genotypes accumulated similar amounts of K⁺, ranging from ∼2.70–2.73 µg and ∼2.68–3.10 µg per seedling, respectively (Figure 5).

### Transcriptome analysis identifies NaCl- and CaCl₂-responsive *PP2C*s modulated by CAMTA6 during Arabidopsis germination

To investigate the extent of PP2C involvement in the germinating seedling response to salt, calcium, and CAMTA6 activity, we analysed two complementary transcriptome datasets. One compared germinating *camta6-5* mutants to wild type seedlings under NaCl stress (Shkolnik et al., 2019; suTable S1), and the other profiled wild type responses to NaCl, CaCl₂, or their combination (Chandran et al., 2024; Table S2). This analysis revealed several *PP2C*s—including *PP2C49*—as prominently salt-responsive in a CAMTA6-dependent manner.

Cross-examination of the transcriptome datasets identified 63 *PP2C* genes, including *PP2C49*, as responsive to the tested conditions (Tables S1 and S2). We identified 40 genes that overlapped in the two transcriptome datasets and organized them into a phylogenetic tree consisting of 9 groups (A–H and J) using the RAxML and FigTree tools (Figure S10), as reviewed by (Schweighofer et al., 2004). Among them, 12 genes were detected in both datasets and met the selection criteria of fold change greater than 1.25 and *P* < 0.05 in at least one of the examined conditions (Table S3). *PP2C49* expression was mildly reduced in the *camta6-5* mutant background, suggesting a potential positive regulatory role for CAMTA6. Nonetheless, treatment with CaCl₂ led to more pronounced downregulation of *PP2C49* (fold change = –3.29; *P* = 0.03; Tables S2 and S3), supporting its involvement in Ca^2+^-signaling pathways, in agreement with our findings (Figure 2).

Among the four group A *PP2C*s identified, *AHG1* and *HAI2* exhibited strong NaCl-inducible expression in the wild type, with fold changes of 9.35 (P = 1.42E⁻⁶) and 14.97 (*P* = 1.65E⁻⁶), respectively (Table S3). In the *camta6-5* mutant, this induction was markedly attenuated, with fold changes reduced to 4.55 (*P* = 2.61E⁻⁵) and 5.39 (*P* = 5.58E⁻⁵), reflecting a 35.74% and 64% loss of NaCl-inducible transcript accumulation, respectively. These findings suggest that CAMTA6 plays a critical role in mediating the salt-induced transcriptional activation of *AHG1* and *HAI2*. In contrast, the other identified group A *PP2Cs*, *ABI1* and *ABI2*, showed only minor responsiveness to NaCl and were largely unaffected by the *camta6-5* mutation (Table S3), indicating a CAMTA6-independent regulatory mechanism.

Of the three group F *PP2C*s identified, the one encoded by AT5G34740 exhibited a prominent NaCl response, with a fold change of 74.75 (*P* = 3.02E-6) in the wild type. This induction was substantially reduced by 77.66%, dropping to 16.7 (*P* = 7.02E-5) in the *camta6-5* mutant (Table S3). These data strongly suggest a role for this PP2C in the CAMTA6-modulated germinating seedling’s response to salt.

These data expand our understanding of how CAMTA6 modulates early responses to salt, in part through selective regulation of *PP2C*s.

## DISCUSSION

### Ca²⁺ modulates the spatial expression of *HKT1;1* via CAMTA6

Seed germination is a tightly controlled developmental process that is highly sensitive to abiotic stresses, including salinity (Nonogaki et al., 2010). Calcium signaling plays a central role in mediating adaptive responses during germination under stress conditions (Steinhorst and Kudla, 2013). In this study, we demonstrate that CAMTA6, a Ca^2+^/calmodulin-dependent transcription factor, modulates seed germination under salt-stress conditions, linking calcium signaling to ionic homeostasis mechanisms.

Our findings reveal that CAMTA6 modulates the expression of HKT1;1, a Na⁺/K^+^ transporter known to limit sodium accumulation in plant tissues (Sunarpi et al., 2005). Under control conditions, CAMTA6 expression is predominantly confined to the cotyledons of germinating Arabidopsis seedlings. Upon salt stress, CAMTA6 expression is markedly enhanced and becomes strongly localized to the cotyledon margins (Figure 1, a and b). In contrast, HKT1;1 expression is broadly detected across all seedling tissues under both control conditions and in response to NaCl treatment, except the cotyledon margins (Figure 1, a and c).

This spatial distinction suggests that CAMTA6 may act as a negative regulator of HKT1;1 expression, particularly at the cotyledon margins, and that its upregulation under salt stress serves to fine-tune HKT1;1 spatial expression in response to that stress. Upon CaCl₂ treatment, CAMTA6 expression is reduced and restricted to the cotyledon margins, whereas HKT1;1 expression becomes predominantly localized in the radicle (Figure 1, a and c). This pattern suggests that elevated Ca²⁺ levels negatively regulate CAMTA6 expression, leading to radicle-confined expression of HKT1;1, similar to its pattern observed in the *camta6* mutant background (Figure 1, a and c).

In the absence of functional CAMTA6, *HKT1;1* expression is restricted to the radicle tissues across all tested conditions, a pattern that correlates with enhanced salt tolerance during seed germination (Shkolnik et al., 2019; Chandran et al., 2024). These observations suggest that *HKT1;1* expression may also be regulated by additional factors, potentially including other CAMTA family members or alternative Ca²⁺-responsive transcriptional regulators capable of recognizing the ABRE sequence (Kaplan et al., 2006; Finkler et al., 2007b). In the *camta6-5* mutant background, *PP2C49* expression was no longer confined to the radicle but extended broadly throughout the emerging seedling under all tested conditions, indicating a loss of spatial regulation. Nonetheless, pronounced expression at the radicle was still evident, as in the wild type. This localized expression (Figure 2a) suggests a role for PP2C49 in environmental sensing and in coordinating the transition from seed coat rupture to post-germination growth. The expanded expression domain in *camta6-5* implies that CAMTA6 may function to spatially constrain *PP2C49* transcription during early seedling development, potentially linking calcium signaling to the spatial control of phosphatase activity.

*camta6* mutants exhibit enhanced tolerance to salinity during germination (Shkolnik et al., 2019), a phenotype that appears to run counter to the observed broad and unregulated expression of *PP2C49* in this background. Given the known inhibitory effect of *PP2C49* on *HKT1;1* function (Chu et al., 2021), such misexpression would be expected to suppress *HKT1;1* activity and increase salt sensitivity. One possible explanation for this discrepancy is that loss of spatial regulation impairs the normal function of *PP2C49*, potentially through mislocalization, disruption of interaction networks, or dosage imbalance. Similar cases have been reported for other group A PP2Cs, including *ABI1*, where constitutive or misregulated expression resulted in dominant-negative effects or altered subcellular dynamics, ultimately compromising phosphatase activity (Merlot et al., 2001; Saez et al., 2004). These observations suggest that precise spatial and temporal control of *PP2C49* expression is essential for its function during germination under saline conditions, and that CAMTA6 contributes to this regulatory specificity. The radicle-confined expression of *HKT1;1* may enhance salt tolerance during germination by limiting Na⁺ accumulation in the cotyledons, where salt-sensitive processes such as chloroplast biogenesis are critical for successful germination and the transition to a photosynthetically active seedling (Peharec-Štefanić et al., 2013). Nevertheless, this explanation should be directly assessed by measuring the Na^+^ concentrations in cotyledons of control and salt-stressed seedlings.

Previous studies have shown that germinating *hkt1* mutants accumulate higher levels of Na⁺ and lower levels of K⁺ under salt stress, compared to wild type seedlings, indicating impaired ion homeostasis (Shkolnik et al., 2019; Chandran et al., 2024). In mature seedlings and adult plants, HKT1;1 is mainly expressed in the vasculature, where it retrieves Na⁺ from the xylem sap to protect the shoot from salt injury (Sunarpi et al., 2005; Møller et al., 2009). By contrast, in germinating seedlings, HKT1;1 expression appears more widespread, including radicle and inner cotyledon tissues, which may facilitate Na⁺ efflux to the medium. Loss of this activity in the *hkt1;1* mutant would thus result in increased Na⁺ accumulation and a pronounced reduction of the K⁺/Na⁺ ratio under salinity stress (0.13 ± 0.05 versus 0.32 ± 0.01 in Col-0, mean ± SD, Figure 5c; Table 1). Importantly, limiting Na⁺ accumulation is essential to maintain K⁺ homeostasis during early development, a balance critical for seedling establishment under salinity stress (Deinlein et al., 2014).

**Table 1.**
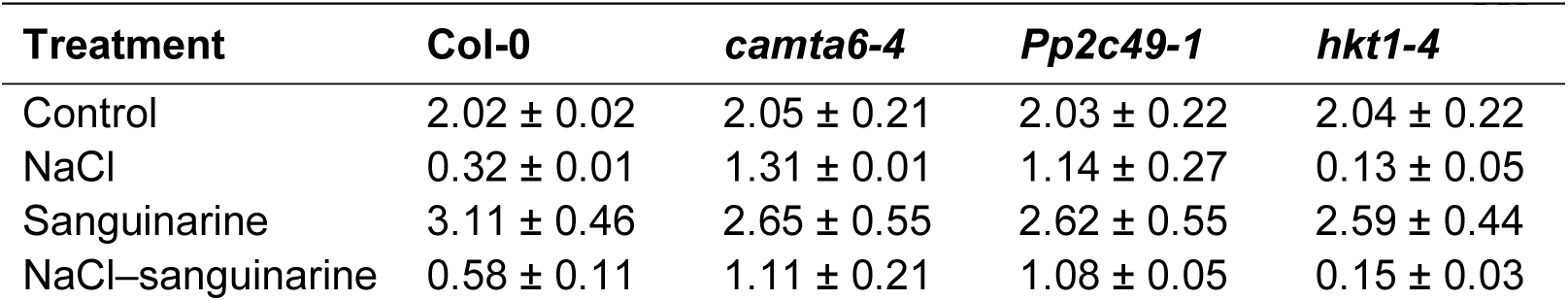
K⁺/Na⁺ ratios in germinating seedlings of *camta6-4*, *pp2c49-1*, and *hkt1-4* mutants under salt and/or sanguinarine treatments. Seedlings were germinated in the presence of 0 (control), 1 µM sanguinarine, 150 mM NaCl, or both, as indicated. Ion contents were determined after 16 h of treatment using ICP-OES, and K⁺/Na⁺ ratios were calculated accordingly. Data represent means ± SD of three biological replicates.

Interestingly, Na⁺ levels did not change significantly in the *camta6-4* and *pp2c49-1* mutants, and this was reflected in their ability to maintain much higher K⁺/Na⁺ ratios under salt stress (1.31 ± 0.01 and 1.14 ± 0.27, respectively, Figure 5c; Table 1). Our model provides a possible explanation for this observation: in germinating seedlings, loss of CAMTA6 or PP2C49 removes their negative regulation of HKT1;1—at the transcriptional and post-translational levels, respectively—resulting in stabilized HKT1;1 expression in the radicle. This compensatory effect helps maintain Na⁺ extrusion to the medium, thereby preventing Na⁺ accumulation and sustaining K⁺ homeostasis. Consistently, under combined NaCl–sanguinarine treatment, the K⁺/Na⁺ ratio in Col-0 recovered only partially (0.58 ± 0.11), whereas *camta6-4* and *pp2c49-1* maintained values close to unity (1.10 ± 0.20 and 1.08 ± 0.05, respectively), in sharp contrast to *hkt1;1*, which remained severely impaired (0.15 ± 0.03, Figure 5c; Table 1).

### Ca^2+^ modulation of seed germination under salt stress through PP2Cs

Our findings suggest that PP2C49 functions as a critical link between CAMTA6 and HKT1;1 within the Ca^2+^-signaling pathway. To further explore this regulatory network, we searched for additional PP2Cs that are responsive to NaCl and Ca²⁺ or exhibit altered expression in the *camta6* mutant background, based on available transcriptome datasets (Shkolnik et al., 2019; Chandran et al., 2024).

Transcriptome analysis revealed that salt stress induces the expression of group A PP2C genes *AHG1* and *HAI2* (Figure S10, Table S3), both of which are well-established negative regulators of ABA signaling (Fujii et al., 2009; Park et al., 2009). Interestingly, the salt-induced expression of *AHG1* and *HAI2* was significantly attenuated in the *camta6-5* mutant—by 35.7% and 64%, respectively (Table S3)—suggesting that CAMTA6 contributes to their transcriptional activation in response to salinity. Given that CAMTAs function as Ca^2+^-regulated transcription factors implicated in abiotic stress responses (Doherty et al., 2009; Galon et al., 2010), this finding raises the possibility that CAMTA6 integrates Ca^2+^- and ABA-signaling pathways under salt stress. Notably, other group A PP2Cs, such as *ABI1* and *ABI2*, showed only modest induction by salt, and their expression was largely unaffected in *camta6-5*, highlighting a selective role for CAMTA6 in modulating a specific subset of ABA-related stress-responsive genes.

Among several group F PP2C-encoding genes identified in our transcriptomic analysis, AT2G34740 exhibited particularly strong induction under salt stress (Table S3), with strong upregulation in both wild type (74.75-fold) and *camta6-5* mutant (16.7-fold) seedlings, suggesting that CAMTA6 may contribute to its transcriptional activation under saline conditions. While group A PP2Cs are well-known negative regulators of ABA signaling (Park et al., 2009; Umezawa et al., 2009) group F members remain less characterized. Nevertheless, functional studies in transgenic tobacco have shown that overexpression of a group F PP2C from wheat can enhance tolerance to salt stress and confer ABA insensitivity (Hu et al., 2015), pointing to roles beyond canonical ABA signaling. The salt-inducible expression of AT2G34740 during germination, coupled with its reduced induction in the *camta6* mutant, identifies it as a promising candidate for further investigation into the contribution of group F PP2Cs to early-stage stress adaptation.

Sanguinarine, a plant-derived benzophenanthridine alkaloid, was found here to enhance salt-stress tolerance during seed germination (Figure 3). Based on its reported selective inhibition of PP2Cs in mammalian systems (Aburai et al., 2010), sanguinarine was applied to test its effect on ABA-mediated inhibition of germination. However, sanguinarine treatment did not alleviate ABA-mediated inhibition of germination in wild type Arabidopsis seeds (Figure S4). Furthermore, mutants of *PP2C49* and *HKT1;1* displayed germination rates comparable to the wild type across a range of ABA concentrations (Figure S5). These results suggest that PP2C49 and HKT1;1 affect germination through pathways that are largely independent of canonical ABA signaling. The combined lack of ABA-sensitivity alteration in these mutants and the ineffectiveness of sanguinarine in modulating ABA’s inhibitory effect imply that the PP2Cs targeted by sanguinarine are unlikely to be major contributors to ABA-mediated germination inhibition in Arabidopsis. While further validation of sanguinarine’s specificity for plant PP2Cs is warranted, these findings underscore the functional specialization within the PP2C family, supporting the view that only a subset of group A PP2Cs—such as AHG1 and HAI2—function as ABA-responsive regulators during salt stress, potentially downstream of CAMTA6. It is important to note that PP2C49 belongs to group G of the PP2C family, distinct from group A members such as PP2CA/PPAC, which are well-established negative regulators of ABA signaling (Park et al., 2009; Umezawa et al., 2009). This classification is consistent with our findings that the salt-related phenotypes of *pp2c49* and *hkt1* mutants are largely ABA-independent (Figure S5).

The lack of differential promoter activity for *CAMTA6*, *PP2C49*, and *HKT1;1*, along with the similar root development observed in the corresponding mutants upon sanguinarine treatment (Figures S7 and S8), suggests that the sanguinarine-promoted tolerance mechanism operates predominantly during germination and is no longer active at later developmental stages.

Phosphatase activity assays provided biochemical support for the inhibitory effect of sanguinarine on Arabidopsis root PP2Cs. The marked reduction in activity observed in wild-type extracts, contrasted with the lower basal activity of *pp2c49-1* extracts and their reduced response to sanguinarine (Figure S9), is consistent with a functional contribution of PP2C49 to the measured activity. These results suggest that sanguinarine inhibits root-derived PP2C activity, and raises the possibility that PP2C49 is among its primary targets. However, given the large number of PP2Cs expressed in Arabidopsis roots (Xue et al., 2008), and the fact that *pp2cg1* seeds still responded to sanguinarine (Figure S6), these data suggest that PP2C49 is likely among the more relevant targets of sanguinarine, although further work is needed to fully define its specificity.

In summary, our results suggest that CAMTA6 mediates responses to salt stress during germination by integrating Ca^2+^ signals into transcriptional regulation, in part through modulation of specific PP2Cs, as CAMTA6 is known to regulate multiple target genes (Shkolnik et al., 2019). This Ca^2+^-dependent pathway functions both alongside and partly independently of canonical ABA signaling.

As climate change is accelerating soil salinization worldwide (Rengasamy, 2010), understanding such early stress-response mechanisms is vital for guiding the development of salt-tolerant crops.

## MATERIALS AND METHODS

### Plant materials, growth and stress assays

Arabidopsis (*Arabidopsis thaliana*) Col-0 plants were used in this research. The following mutants were obtained from the Arabidopsis Stock Center in Columbus, OH: *hkt1* (CS68521), *hkt1-4* (CS68092), *pp2c49-1* (SALK_111549C), *pp2c49-2* (SALK_015078C), *pp2cg1-1* (SALK_036544C), *pp2cg1-2* (SALK_200850C). Plants expressing *HKT1;1pro*:GUS, *CAMTA6pro*:GUS, and *PP2C49pro*:GUS were created as described previously (Shkolnik et al., 2019; Chandran et al., 2024).

Seed-surface sterilization and germination assays in the presence of different chemicals were performed as previously described (Shkolnik and Bar-Zvi, 2008). Germination assays on NaCl, CaCl_2_, sanguinarine (sanguinarine chloride hydrate; Sigma-Aldrich) or their combination were performed in the same manner. A seedling was considered germinated when green fully unfurled cotyledons were observed. Treatments with NaCl prior to GUS staining (see GUS staining section further on), and RT-qPCR were performed by seed imbibition at 4 °C in the dark for 2–3 days prior to plating on Whatman filter paper (grade 1) soaked with 0.25X MS medium (Murashige and Skoog, 1962) and supplemented with the indicated chemicals. Pre-germinating seedlings were gently ejected manually, using curved fine forceps, 16 h after starting the treatment. Separation of cotyledon and radicle tissues for RNA isolation and subsequent RT-qPCR analysis was performed by relatively rough seedling ejection from the seed coat and tissue collection in separate test tubes using a fine pipette under a binocular. Treatment of several-day-old seedlings with NaCl, CaCl_2_, sanguinarine, or the indicated combinations was performed by placing the seedlings such that the root was in contact with the agar-solidified 0.25X MS medium supplemented with the indicated chemicals, and the shoot was in the air to avoid direct contact with NaCl as previously described (Shkolnik-Inbar and Bar-Zvi, 2012; Shkolnik et al., 2019). The seedlings were grown vertically in growth chambers under a 16-h light/8-h dark photoperiod at 20 °C.

### GUS staining

GUS staining was performed as previously described (Weigel and Glazebrook, 2002). Briefly, seedlings were treated with 300 μL of 90% (v/v) acetone for 1 min, washed with 1 mL staining solution (50 mM sodium phosphate buffer pH 7.0, 2 mM ferricyanide, 2 mM ferrocyanide, and 0.2% v/v Triton X-100) followed by 1 h incubation at 37 °C in staining solution supplemented with 2 mM X-Gluc (5-bromo-4-chloro-3-indolyl-beta-D-glucuronic acid, cyclohexylammonium salt). Images were taken using the Nikon SMZ18 stereoscope system equipped with a DS-Fi3 camera.

### Quantitative GUS assay

GUS activity was quantified spectrophotometrically as described previously (Jefferson, 1987). Briefly, soluble protein preparations were extracted from 100 mg of GUS-expressing Arabidopsis seedlings using GUS extraction buffer containing 50 mM NaPO_4_ pH 7.0, 10 mM beta-mercaptoethanol, 10 mM Na_2_EDTA, 0.1% v/v- sodium lauryl sarcosine and 0.1% Triton X-100. Reaction was initiated by mixing 50 µL of protein extract in a 0.5 mL total volume of 1 mM of the GUS substrate 4-nitrophenyl β-D-glucuronide (PNPG). The reaction was incubated at 37 °C for 9 h and absorbance was measured at 415 nm using a spectrophotometer.

### RT-qPCR analysis

Total RNA was isolated from pre-germinating seedlings using the ZR Plant RNA MiniPrep Kit (Zymo Research), and total cDNA was synthesized using the High-Capacity cDNA Reverse Transcription Kit (Thermo Fisher Scientific). The reaction mixture was prepared according to the manufacturer’s instructions, with random primers, and supplemented with 1 µg of total RNA. PCR mixtures (10 µL), containing 5 µL Fast SYBR Green Master Mix (Applied Biosystems by Thermo Fisher Scientific), 500 nM reverse and forward primers designed to amplify 80–150 bp of the genes of interest (Table S4), and 20 ng cDNA were subjected to the Rotor-Gene Q 5-Plex HRM real-time PCR system (Qiagen) using the default program. The *PP2A* (AT1G69960) gene was used as an endogenous control. Relative quantification data were analyzed by the Rotor-Gene Q Pure Detection software V2.3.5 (Qiagen).

### Ion-content analysis

Pre-germinating seedlings subjected to control, NaCl, sanguinarine, or combined NaCl–sanguinarine treatments were collected following ejection from seeds. Sample preparation for Na⁺ and K⁺ quantification was performed as previously described (Kalifa et al., 2004; Shkolnik-Inbar et al., 2013). In each biological replicate, 40 seedlings per line were sampled. Ion concentrations were measured using an Arcos ICP-OES spectrometer (Spectro/Ametek).

### Transient promoter activation assay

Promoter fragments of approximately 2 kb (*CAMTA6*, *HKT1;1*, and *PP2CG1*) were amplified using gene-specific primers listed in Supplemental Table S4 and subcloned individually into the pCAMBIA1391Z vector upstream of the GUS reporter gene, with the *CAMTA6* and *HKT1;1* promoters inserted using the KpnI/SalI restriction sites and the PP2CG1 promoter inserted using the PstI/BamHI restriction sites. The *CAMTA6* coding sequence fragment was subcloned into the pBTEX vector downstream of the constitutive CaMV 35S promoter using the KpnI/EcoRI restriction sites to generate the 35S:CAMTA6 (CAMTA6-OE) construct. Each promoter::GUS fusion construct, either alone (control) or co-infiltrated with the CAMTA6-OE construct, was introduced into *Nicotiana benthamiana* leaves via Agrobacterium-mediated infiltration, after which leaf discs were collected from the infiltrated regions and subjected to GUS staining. The empty pCAMBIA1391Z vector served as an additional control. The procedure was repeated at least 10 times independently.

### Protein extraction and phosphatase activity assay

Crude protein extracts were prepared from roots of 4-day-old Arabidopsis seedlings (Col-0 and *pp2c49-1*). Root tissue was harvested, flash-frozen in liquid nitrogen, and ground to a fine powder. Total soluble protein was extracted in ice-cold buffer containing 50 mM Tris-HCl (pH 7.5), 150 mM NaCl, 0.1% (v/v) Triton X-100, and 1 mM dithiothreitol (DTT), supplemented with a protease inhibitor cocktail (Roche). Homogenates were centrifuged at 14,000 × g for 15 min at 4 °C, and the clarified supernatant was collected. Protein concentration was determined by the Bradford assay using bovine serum albumin as a standard (Bradford, 1976).

Phosphatase activity was assayed essentially as described by Hurley et al. (2010) with minor modifications. Reaction mixtures (100 µL) contained 10 µg of total protein and 10 mM p-nitrophenyl phosphate (pNPP; Sigma) in assay buffer (50 mM Tris-HCl pH 7.5, 150 mM NaCl, 5 mM MgCl₂, 0.1% Triton X-100, and 1 mM DTT), in the presence or absence of 1 µM sanguinarine. Reactions were incubated at 30 °C for 30 min and terminated by the addition of 100 µL 0.5 M NaOH. The release of p-nitrophenol from pNPP dephosphorylation was quantified by measuring absorbance at 405 nm using a microplate reader. Control reactions without protein were used to correct for non-enzymatic substrate hydrolysis, and background values were subtracted from all samples. Technical replicates were averaged, and activities were normalized to the mean absorbance of Col-0 controls, which was set to 1.0.

### Phylogenetic analysis

Gene sequences were retrieved from the TAIR10 database (https://www.arabidopsis.org/). Multiple sequence alignment was performed using MAFFT (v7.450) with default parameters. A maximum-likelihood phylogenetic tree was constructed using RAxML (v8.2.12; https://cme.h-its.org/exelixis/web/software/raxml/). The tree was visualized using FigTree (v1.4.4; http://tree.bio.ed.ac.uk/software/figtree/).

### Statistical analysis

Data were analyzed using OriginPro 2024 (OriginLab Corporation) and Microsoft Excel 2021 with the Analysis ToolPak add-in.

## Accession numbers

Accession numbers of the major genes investigated in this research are: *CAMTA6*, AT3G16940; *PP2C49*, AT3G62260; *HKT1;1*, AT4G10310.

## Data Statement

No new transcriptomic datasets were generated in this study. The transcriptome data analyzed here were previously reported in Shkolnik *et al*. (2019) and Chandran *et al*. (2024). All data supporting the findings of this study are included within the article and its Supplementary Information. Additional materials and datasets are available from the corresponding author upon reasonable request.

## Author contributions

Y.K. and D.S. designed the experiments and research plan; Y.K., A.E.J.C., G.S., O.C., S.A.-P., Y.W. and D.S. performed the experiments; Y.K. and D.S. analyzed the data and wrote the article.

## Acknowledgments

This research was supported by the Israel Science Foundation (grant no. 756/20 to D.S.).

## Supplementary material

**Figure S1.**
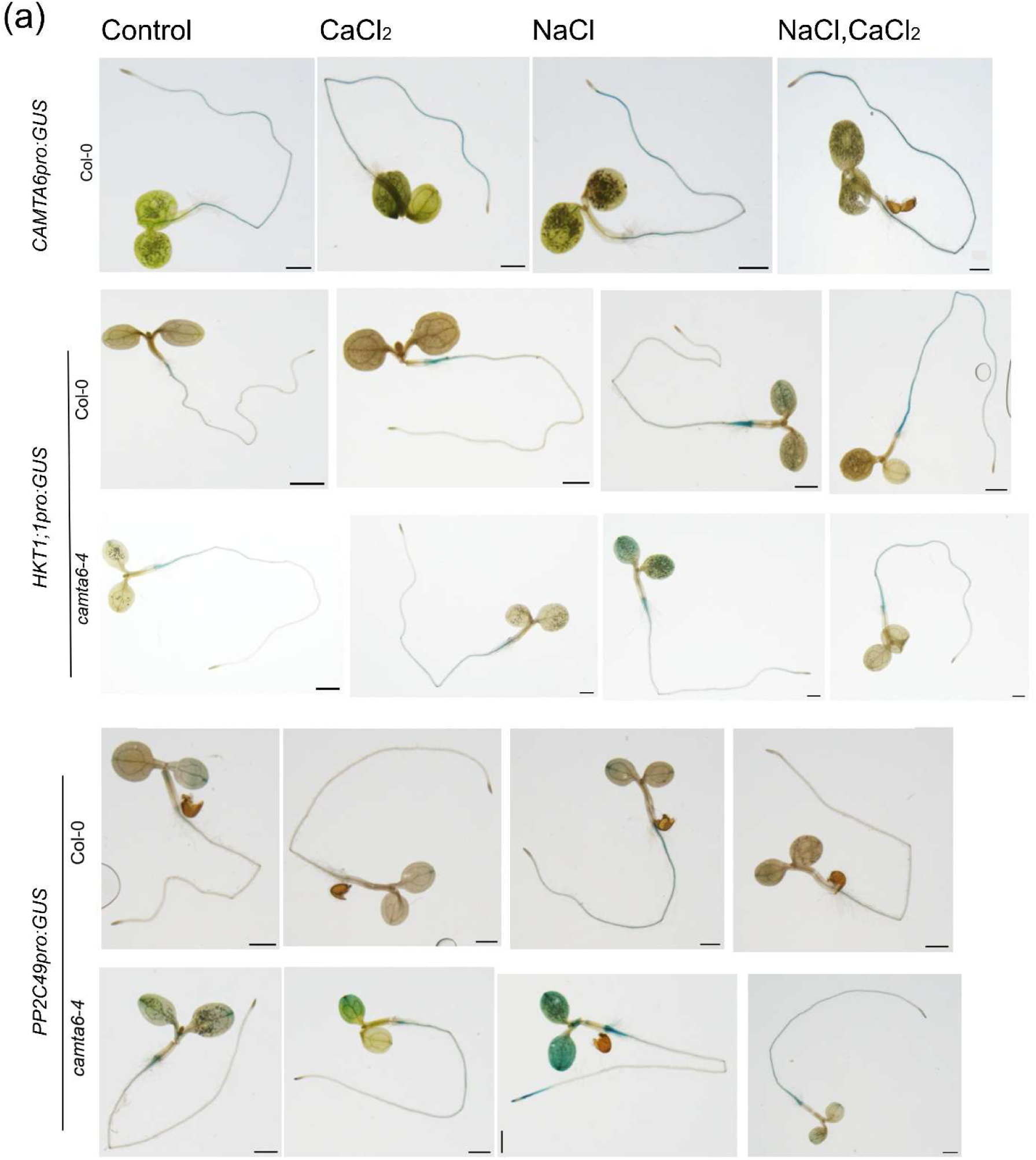
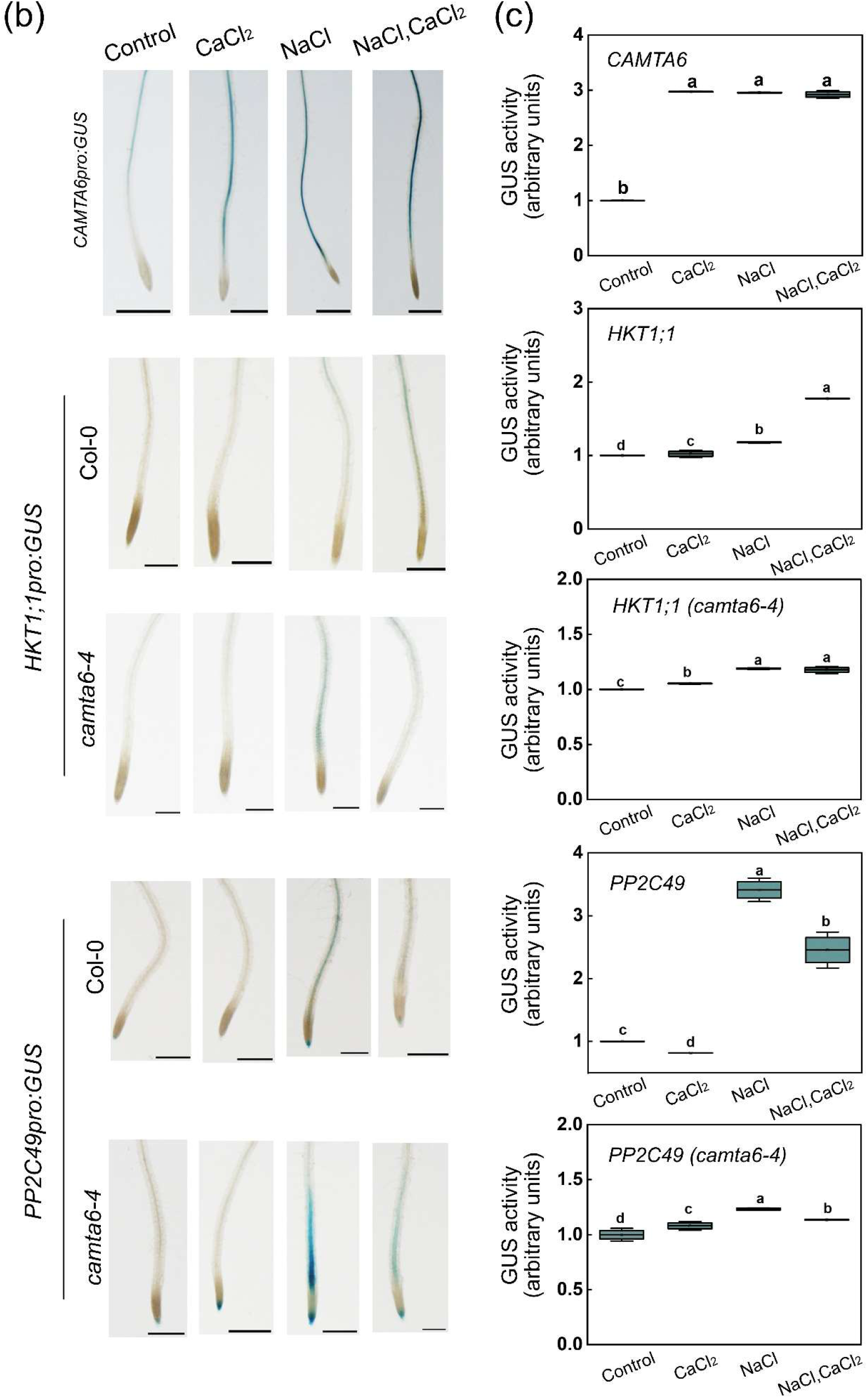
Promoter activity of *CAMTA6*, *HKT1;1*, and *PP2C49* in Arabidopsis seedlings treated with NaCl and/or CaCl₂. (a, b) Five-day-old seedlings of the indicated genotypes were treated with the specified chemicals (NaCl, 150 mM; CaCl₂, 10 mM), GUS-stained (see Materials and Methods), and imaged using a stereomicroscope system. Whole seedlings are shown in (a) and young primary root zones in (b) (bars = 0.1 mm for a; 0.3 mm for b). (c) Spectrophotometric GUS assay. Relative GUS activity was quantified using PNPG as a substrate as described previously (Jefferson, 1987). Bold lines in each box indicate the mean, and whiskers represent ± SE (n = 3 independent biological experiments). Top and bottom sides of the boxes correspond to the third and first quartiles, respectively. Different lowercase letters indicate significantly different values by Tukey’s HSD post hoc test (P < 0.005).

**Figure S2.**
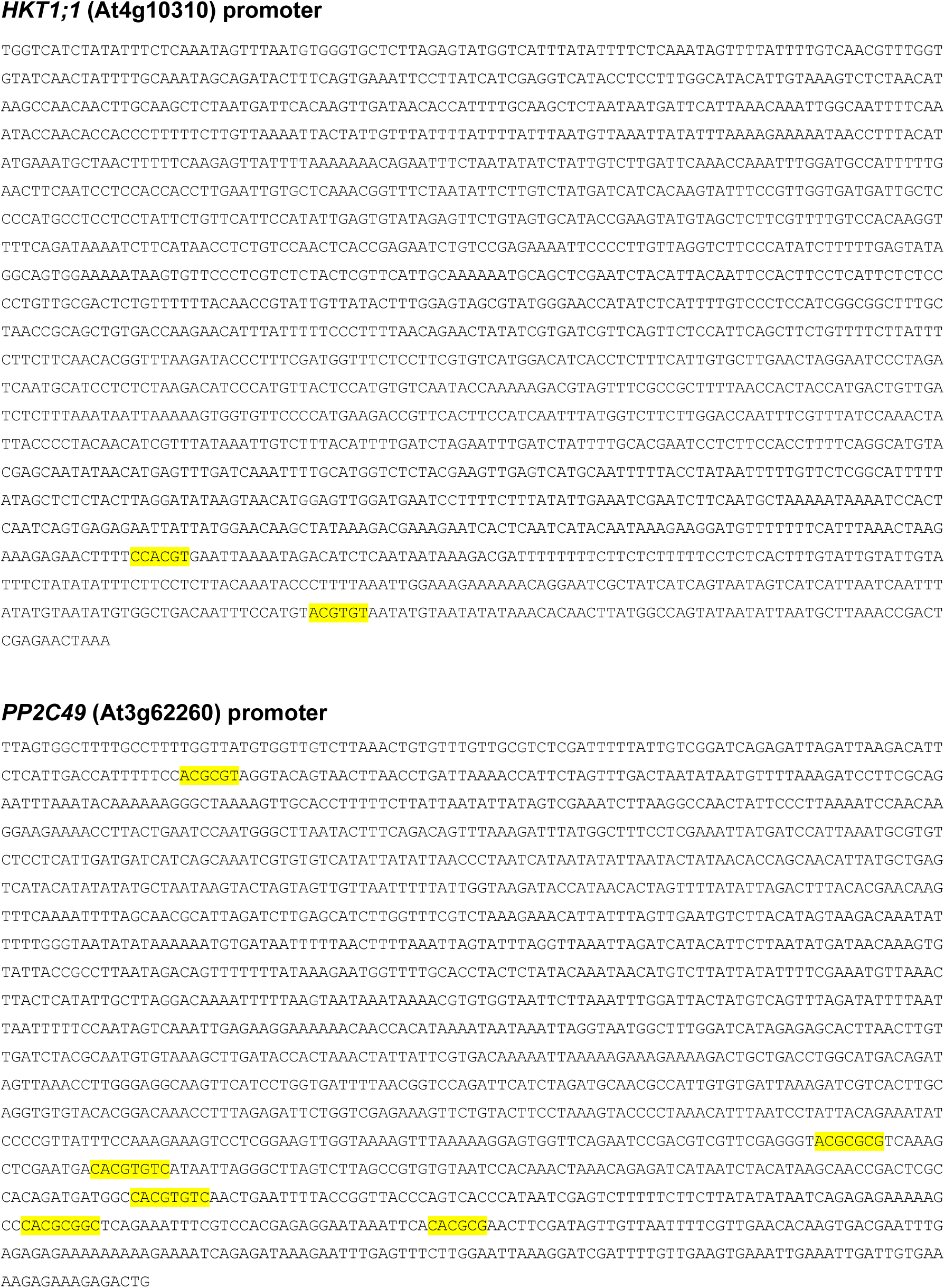
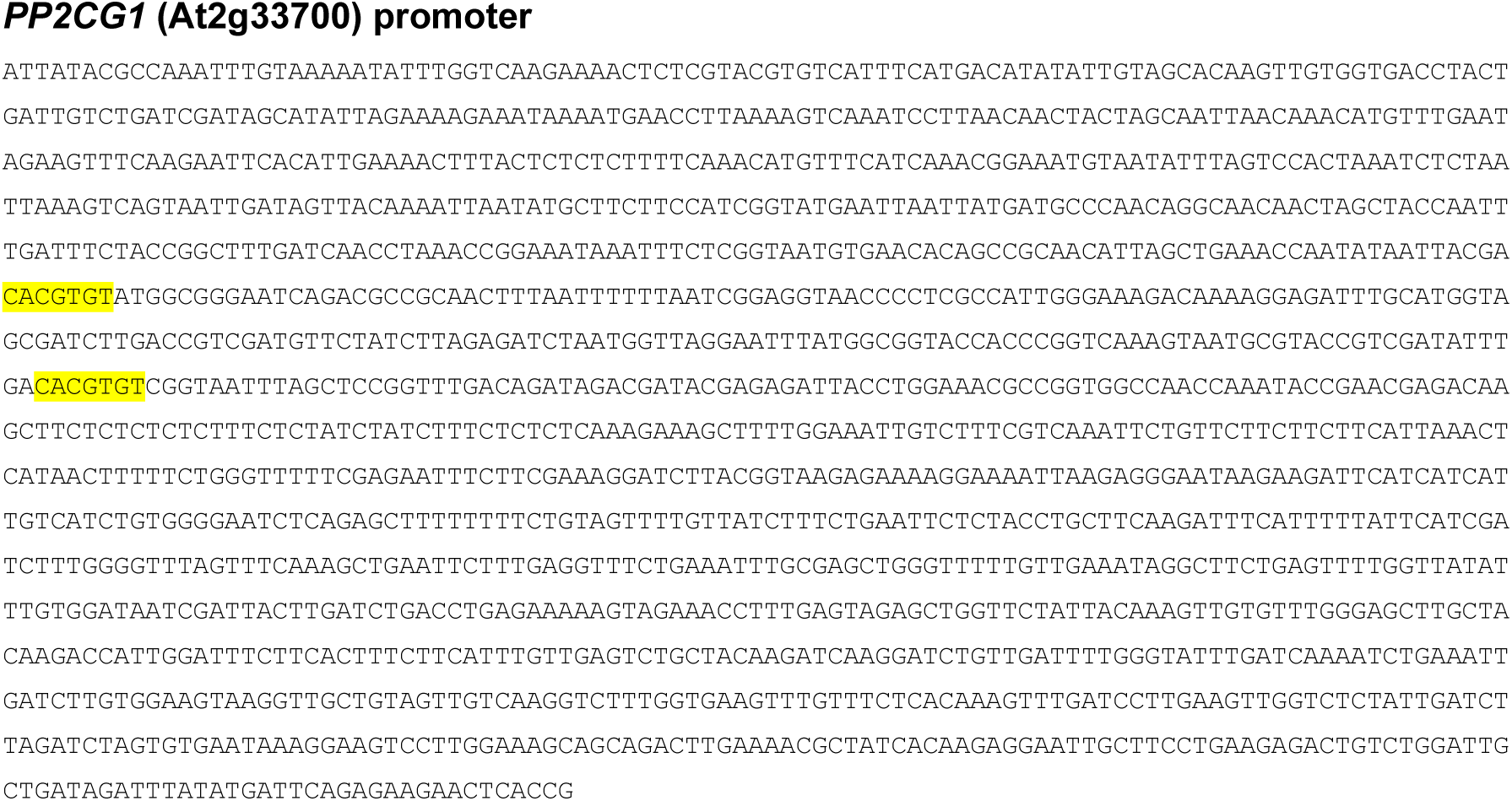
Potential CAMTA-binding elements in the promoters of HKT1;1, PP2C49, and PP2CG1. Promoter sequences were obtained from the TAIR database (Arabidopsis.org). The ABRE/CAMTA-binding element (CACGTGTC) and its coupling element ([C/A]ACGCG[T/C/G]) are highlighted.

**Figure S3.**
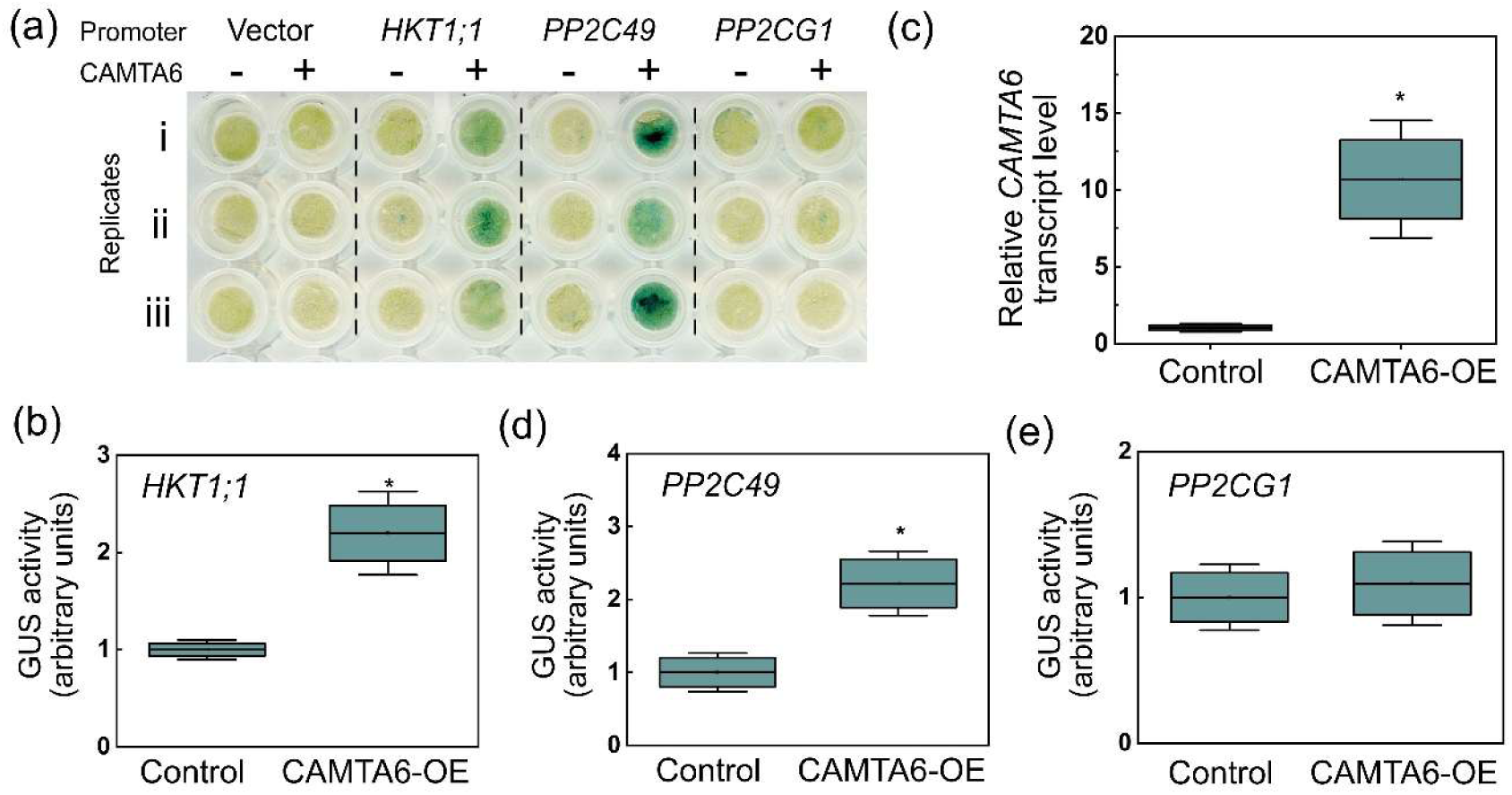
Transient activation of gene promoters by CAMTA6 in *Nicotiana benthamiana* leaves. (a) Gene promoters were subcloned upstream of the GUS reporter in the pCAMBIA1391Z vector. The CAMTA6 coding sequence was cloned downstream of the constitutive CaMV35S promoter (35S:CAMTA6) for overexpression (CAMTA6-OE). Each GUS-fused promoter vector, with or without CAMTA6-OE as indicated, was co-infiltrated into *N. benthamiana* leaves via Agrobacterium, followed by GUS staining of leaf discs from the infiltration region. Empty pCAMBIA1391Z vector served as control. The procedure was performed at least 10 times, and three representative replicates are shown. (b) RT-qPCR quantification of CAMTA6 transcript levels in CAMTA6-OE leaf discs. (c–f) Spectrophotometric GUS assay. Relative GUS activity was quantified using PNPG as a substrate (Jefferson, 1987) and normalized to the control (set as 1). Bold lines in each box represent the means ± SE (three biological experiments), and whiskers indicate ± SE. Top and bottom sides of the boxes correspond to the third and first quartiles, respectively. Asterisks indicate significant differences compared to the control by Student’s t-test (*P* < 0.005).

**Figure S4.**
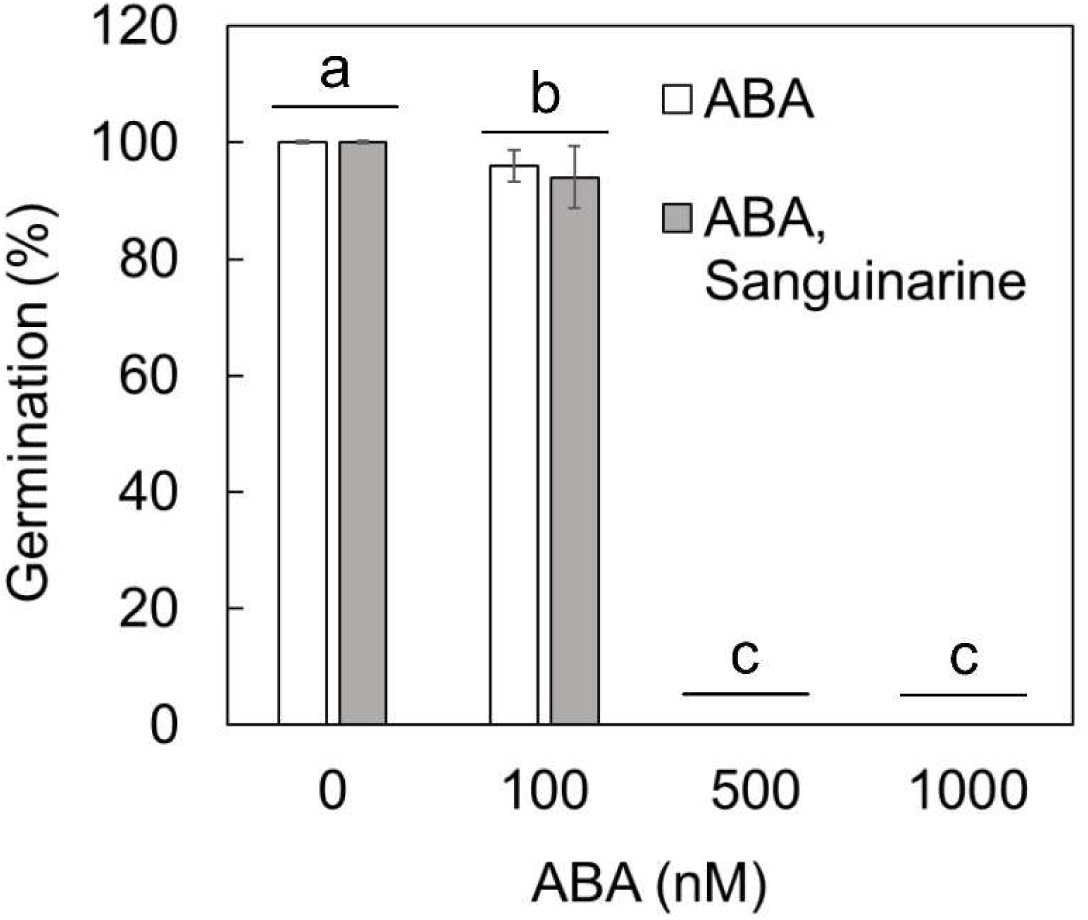
Sanguinarine does not affect ABA-mediated inhibition of Arabidopsis seed germination. Wild-type (Col-0) seeds were sown on agar-solidified 0.25X MS medium supplemented with ABA (0, 100, 500, or 1000 nM) alone or in combination with sanguinarine (1 µM), as indicated. Germination rates were scored 5 days after plating. Data are presented as means ± SD from three independent biological replicates (∼50 seeds per replicate). Different lowercase letters indicate significant differences by Tukey’s HSD post hoc test (*P* < 0.001).

**Figure S5.**
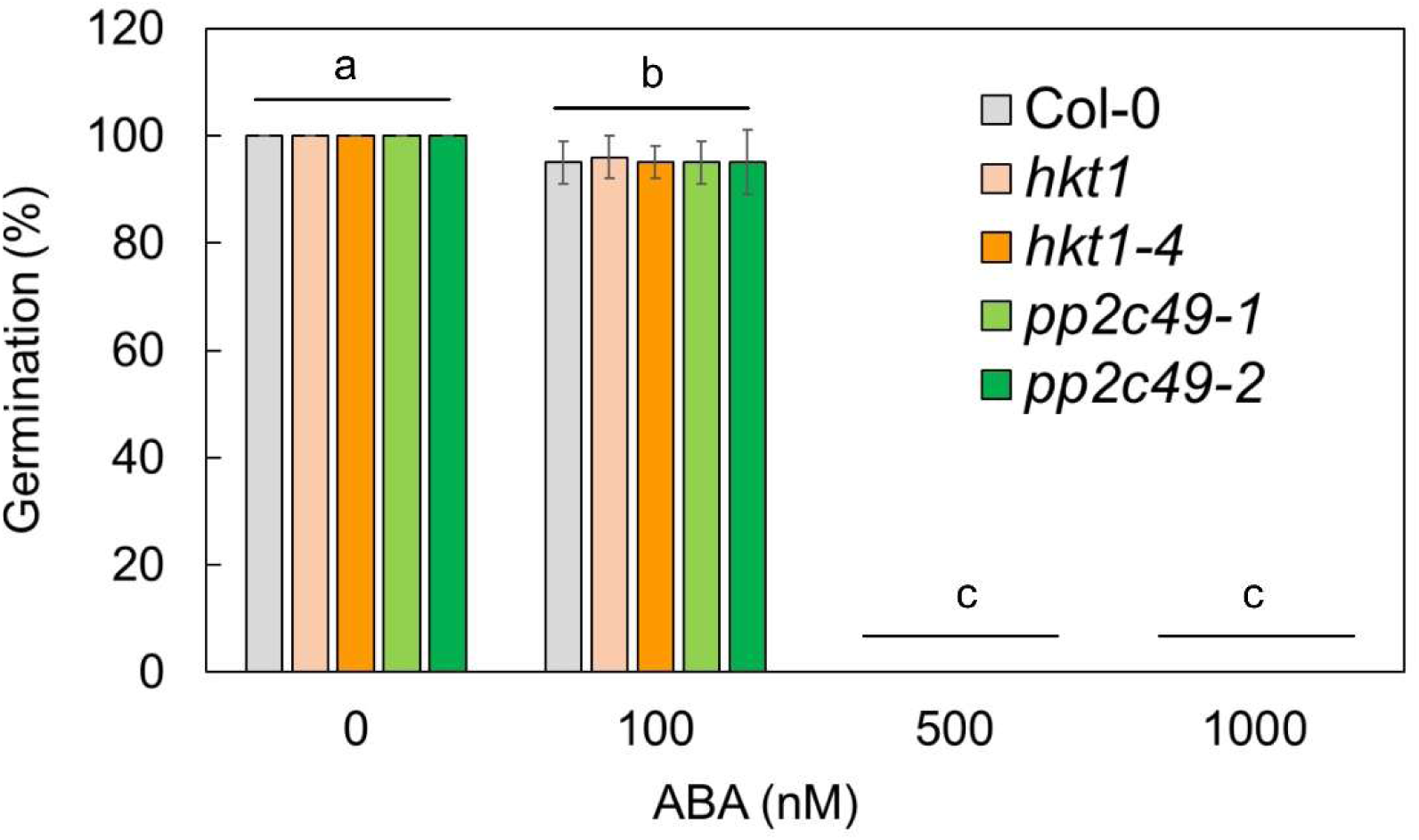
ABA inhibits germination in wild-type and mutant Arabidopsis seeds. Seeds of wild type (Col-0) and *hkt1*, *hkt1-4*, *pp2c49-1*, and *pp2c49-2* mutants were sown on agar-solidified 0.25X MS medium containing the indicated ABA concentrations. Germination was scored 5 days after plating. Data are presented as means ± SD from three independent biological replicates (∼50 seeds per replicate). Different lowercase letters indicate significant differences by Tukey’s HSD post hoc test (*P* < 0.01).

**Figure S6.**
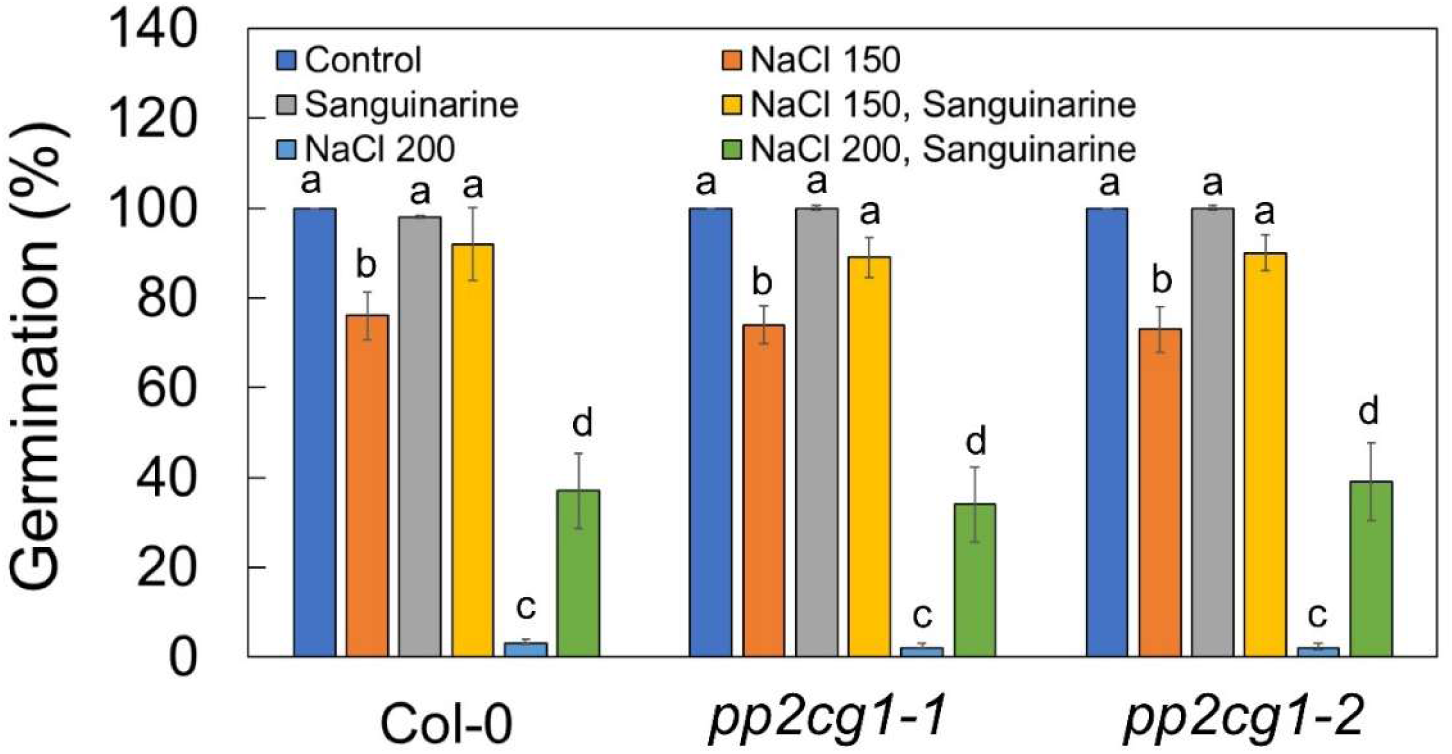
Effects of sanguinarine and NaCl on germination of wild-type and *pp2cg1* mutant Arabidopsis seeds. Seeds of wild-type (Col-0) and *pp2cg1-1* and *pp2cg1-2* mutant alleles were sown on agar-solidified 0.25X MS medium under control conditions or supplemented with the indicated treatments: 1 µM sanguinarine (Sang.), 150 mM NaCl, 200 mM NaCl, Sang. + 150 mM NaCl, or Sang. + 200 mM NaCl. Germination rates were scored 5 days after plating. Data are presented as means ± SD from three independent biological replicates (∼50 seeds per replicate). Different lowercase letters indicate significant differences by Tukey’s HSD post hoc test (P < 0.01).

**Figure S7.**
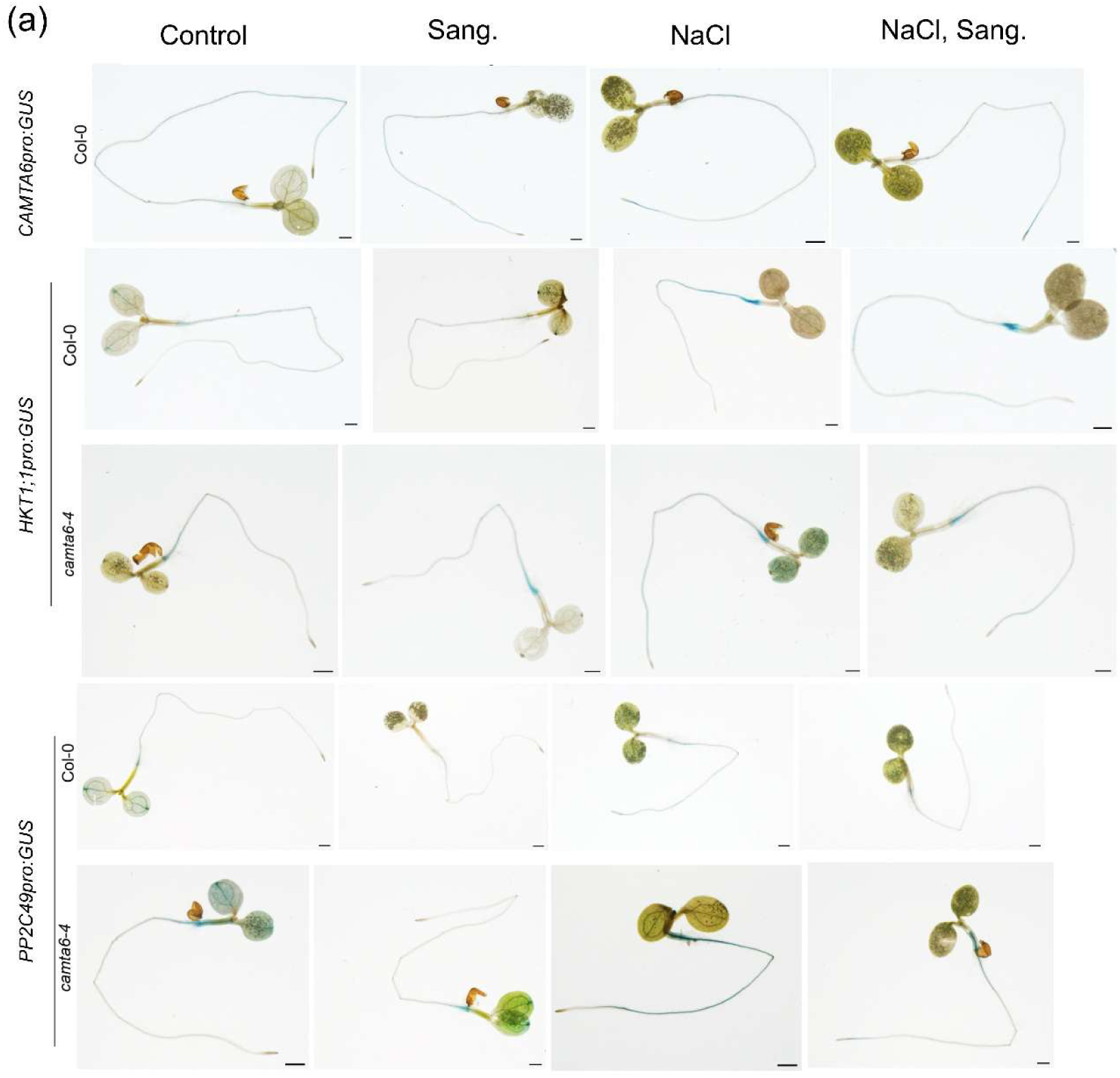
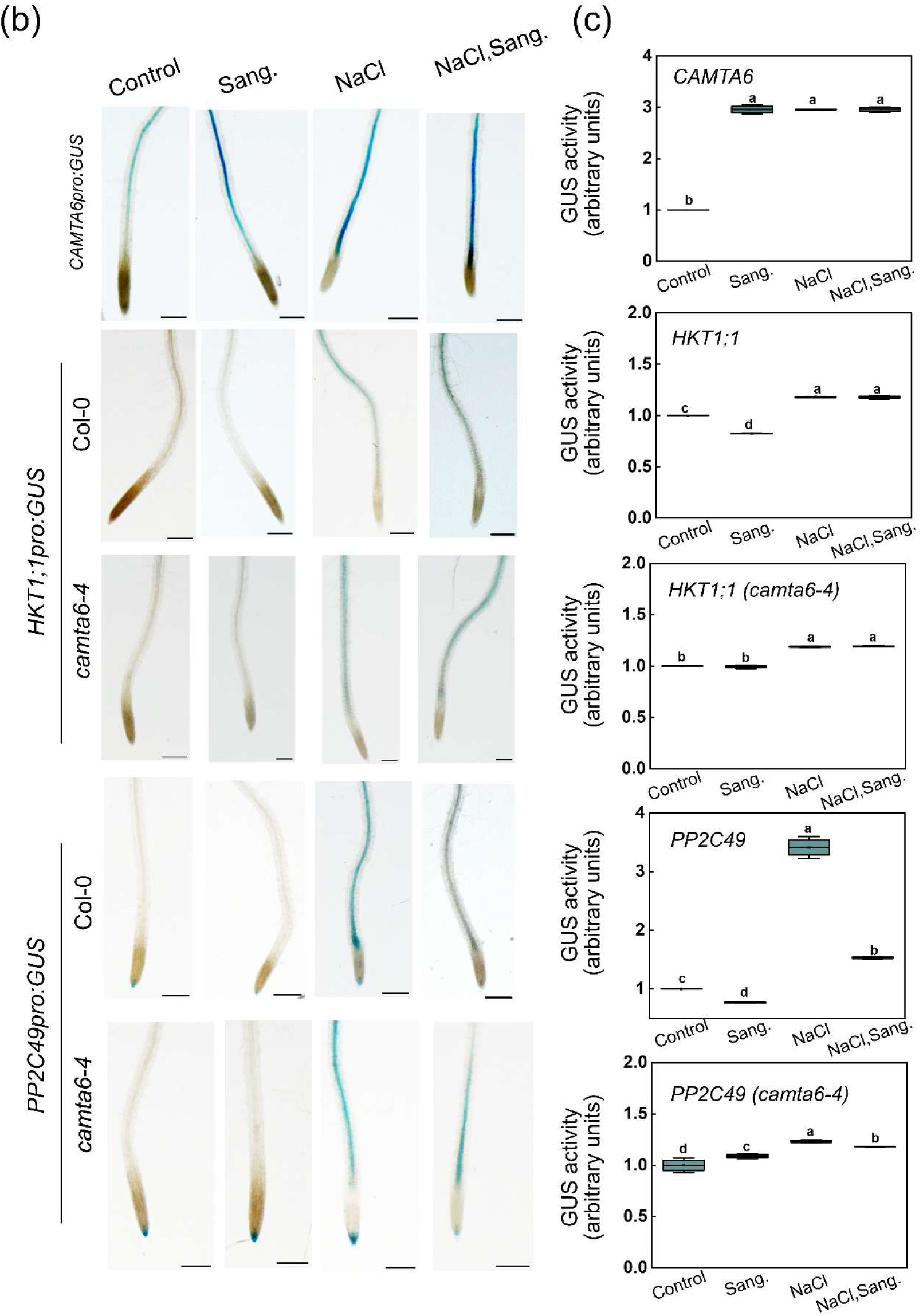
*CAMTA6*, *HKT1;1*, and *PP2C49* promoter activity in Arabidopsis seedlings treated with NaCl and/or sanguinarine (Sang.). (a, b) Five-day-old seedlings of the indicated genotypes were treated with the specified chemicals (NaCl, 150 mM; Sang., 1 µM), GUS-stained (see Materials and Methods), and imaged using a stereomicroscope system. Whole seedlings are shown in (a) and young primary root zones in (b) (bars = 0.1 mm for a; 0.3 mm for b). (c) Spectrophotometric GUS assay. Relative GUS activity of whole root tissue extracts was quantified using PNPG as a substrate (Jefferson, 1987). Bold lines in each box indicate the mean, and whiskers represent ± SE values (n = 3 independent biological experiments). Top and bottom sides of the boxes correspond to the third and first quartiles. Different lowercase letters indicate significantly different values by Tukey’s HSD post hoc test (*P* < 0.005).

**Figure S8.**
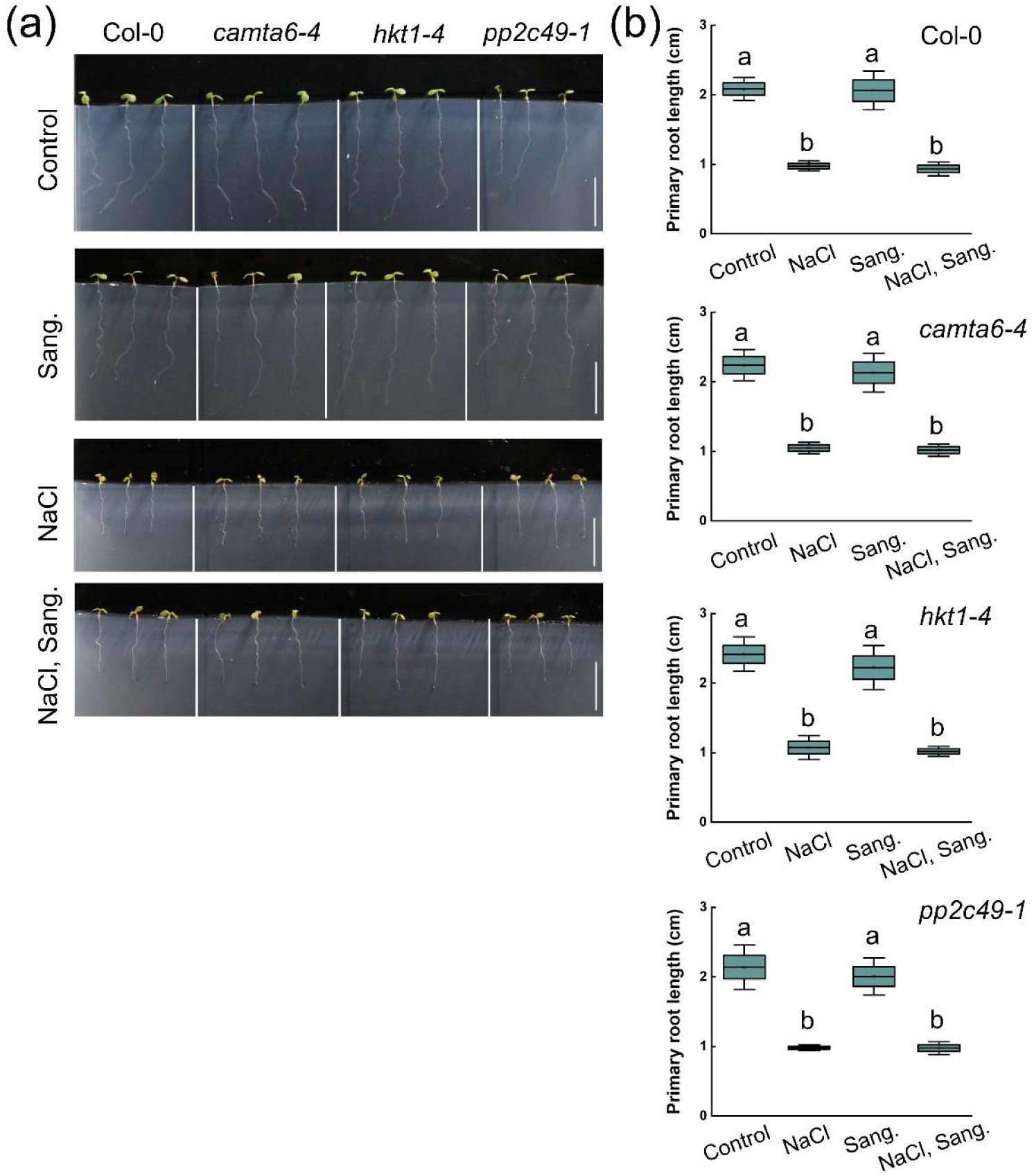
Sanguinarine does not alter primary root growth of Arabidopsis seedlings under control or salt-stress conditions. (a) Four-day-old seedlings of the indicated genotypes were germinated and grown on control medium, then transferred to NaCl (150 mM), Sang. (1 µM), or both treatments for an additional 2 days. Bars = 1 cm. (b) Quantification of primary root length of seedlings treated as in (a). Bold lines in each box represent the means ± SD (three independent biological experiments, 10 seedlings each). Top and bottom sides of the boxes correspond to the third and first quartiles of the distribution. Different lowercase letters indicate significantly different values by Tukey’s HSD post hoc test (*P* < 0.001).

**Figure S9.**
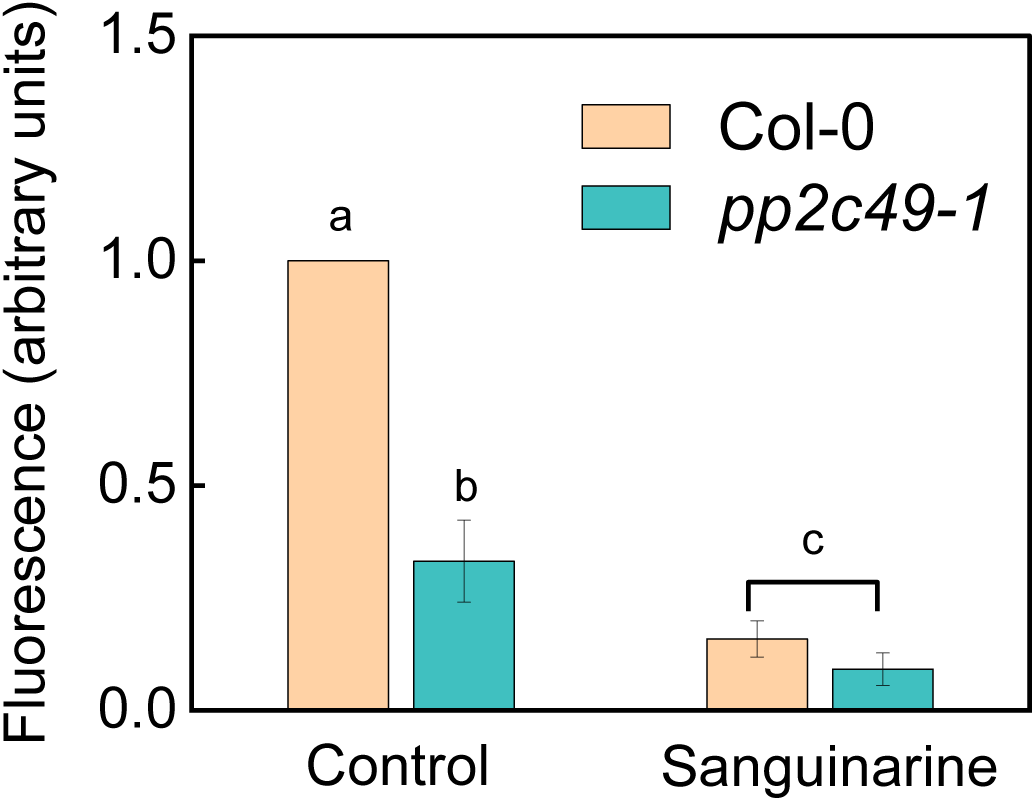
Phosphatase activity in roots of wild-type and *pp2c49-1* Arabidopsis seedlings in the presence or absence of sanguinarine. Crude protein extracts were prepared from roots of 4-day-old Col-0 and pp2c49-1 seedlings. Phosphatase activity was measured using the fluorogenic substrate pNPP. Reactions contained 10 µg of total protein and were incubated with 10 mM pNPP at 30 °C for 30 min, with or without 1 µM sanguinarine. The release of p-nitrophenol was quantified by absorbance at 405 nm. Data are presented as means ± SD from three independent biological replicates, each with three technical replicates. Activity values were normalized to the Col-0 control (set to 1.0). Different lowercase letters indicate significant differences according to Tukey’s HSD post hoc test (*P* < 0.01).

**Figure S10.**
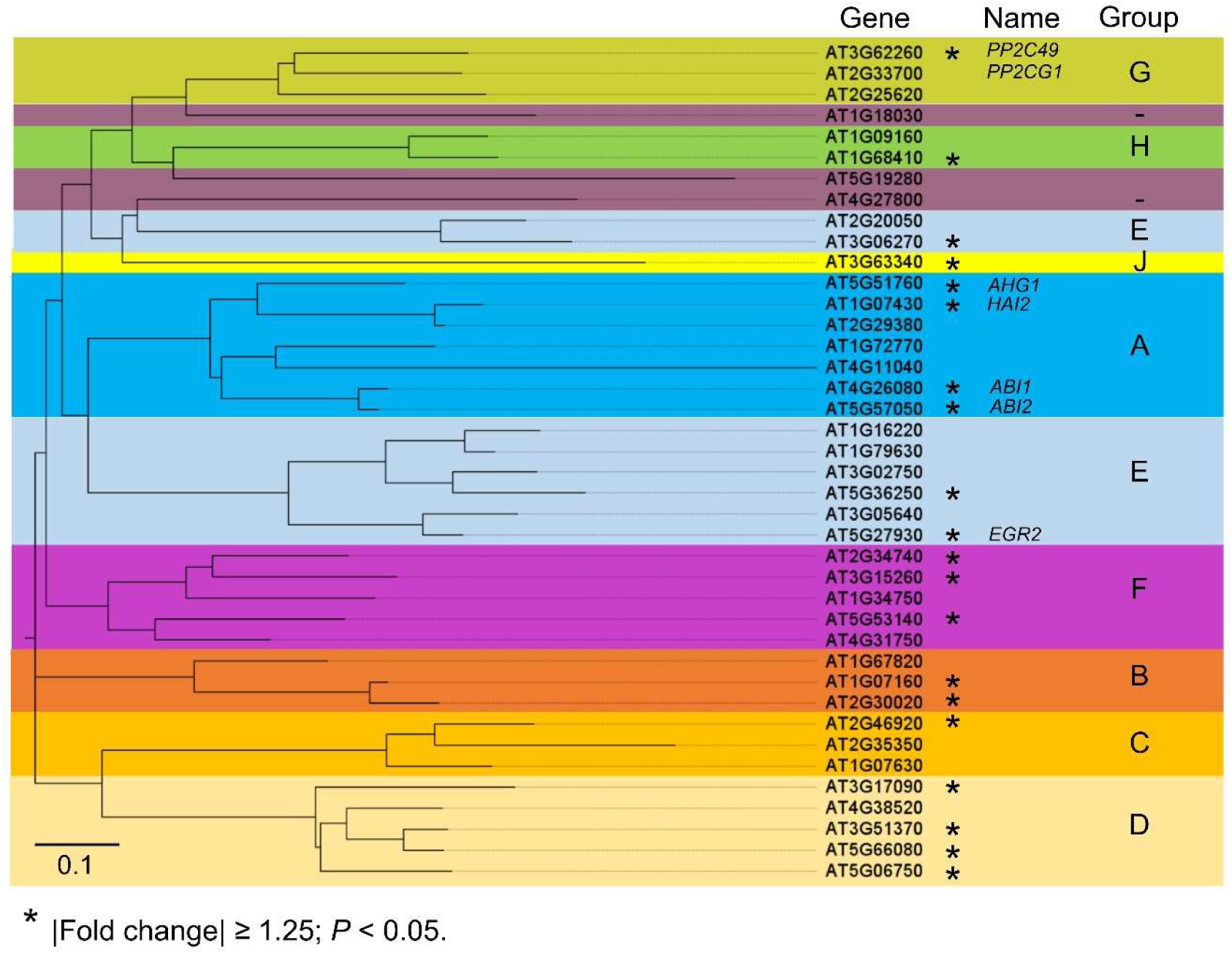
Phylogenetic classification of 40 overlapping PP2C genes identified in transcriptome datasets. Genes were grouped into A–H and J based on maximum-likelihood phylogenetic analysis using RAxML and visualized with FigTree. Twelve genes meeting the selection criteria of absolute fold change ≥ 1.25 and *P* < 0.05 in at least one condition are indicated with an asterisk (*). Gene names and statistical values are provided in Supplementary Tables S1–S3. Scale bar represents 0.1 substitutions per site.

## Supporting table legends

**Table S1.** List of differentially expressed *PP2C* genes identified in comparisons between wild type *Arabidopsis* (Col-0) and the *camta6-5* mutant under control and NaCl-treated conditions, in germinating seedlings. Genes that met the selection criteria of absolute fold change ≥ 1.25 and *P* < 0.05 were considered differentially expressed. The following pairwise comparisons are shown: (1) *camta6-5* NaCl vs. *camta6-5* control; (2) Col-0 NaCl vs. Col-0 control; (3) *camta6-5* control vs. Col-0 control; and (4) *camta6-5* NaCl vs. Col-0 NaCl. Data were extracted from Shkolnik et al. (2019).

**Table S2.** List of differentially expressed *PP2C* genes upregulated and downregulated in response to CaCl₂, NaCl, and their combination, compared to untreated controls, in germinating seedlings of wild type *Arabidopsis* (Col-0). Genes that met the selection criteria of absolute fold change ≥ 1.25 and *P* < 0.05 were considered differentially expressed. The following pairwise comparisons are shown: (1) CaCl₂ vs. control; (2) NaCl vs. control; (3) NaCl–CaCl₂ vs. control; (4) NaCl vs. CaCl₂; (5) NaCl–CaCl₂ vs. CaCl₂; and (6) NaCl–CaCl₂ vs. NaCl. Data were extracted from Chandran et al. (2023).

**Table S3.** List of differentially expressed *PP2C* genes shared between two independent datasets (Shkolnik et al., 2019; Chandran et al., 2023). Genes were included if they met the selection criteria of absolute fold change ≥ 1.25 and *P* < 0.05 in at least one comparison within each dataset. The table presents *PP2C* genes differentially expressed in germinating seedlings of wild type *Arabidopsis* (Col-0) in response to control, NaCl, CaCl₂, and combined NaCl-CaCl₂ treatments, and in the *camta6-5* mutant under control and NaCl-treated conditions.

**Table S4.** List of primers used in this study.

